# Pervasive phosphorylation by phage T7 kinase disarms bacterial defenses

**DOI:** 10.1101/2024.12.20.629319

**Authors:** Tara Bartolec, Karin Mitosch, Clément Potel, Federico Corona, Alessio Ling Jie Yang, Mira Lea Burtscher, Alexandra Koumoutsi, Isabelle Becher, Jacob Bobonis, Nicolai Karcher, Marco Galardini, Athanasios Typas, Mikhail M. Savitski

## Abstract

Bacteria and bacteriophages are in a constant arms race to develop bacterial defense and phage counter-defense systems. Currently known phage counter-defense systems are specific to (the activity of) the targeted bacterial defense system. Here, we uncover a mechanism by which the T7 bacteriophage broadly counteracts bacterial defenses using protein phosphorylation. We show that the T7 protein kinase (T7K), which was believed to specifically redirect the function of a few host proteins, is in fact a hyper-promiscuous, dual-specificity kinase enacting a massive wave of phosphorylation on virtually all host and phage proteins during infection. The scale of phosphorylation vastly exceeds the number of previously known phosphorylation events in *E. coli,* has no sequence motif specificity, and results in a higher proteome-wide phosphorylation density than that of mammalian cells which encode ∼ 500 kinases. Stoichiometry analysis of phosphorylation sites revealed a strong bias of T7K activity towards nucleic acid-binding substrates, which we show is mediated by its C-terminal DNA-binding domain. This specificity for highly stoichiometric phosphorylation of nucleic acid-binding proteins enables the deactivation of DNA-targeting or - containing bacterial defense systems. We provide mechanistic insight into how T7K weakens two such defense systems, Retron-Eco9 and DarTG1, through specific phosphorylation events, with single phosphomimetic mutations in key sites of the toxins abolishing defense. Finally, by screening a large collection of *E. coli* strains, we provide evidence of broad counter-defense capacities for T7K in nature, as strains counteracted contain diverse bacterial defense systems.

## Introduction

Bacteria and their viruses (bacteriophages, or simply phages) engage in an intricate evolutionary arms race, and employ sophisticated molecular strategies to overcome each other’s defenses^1^. In the recent past, our understanding of the bacterial immune system repertoire and its underlying mechanisms^2^ have increased dramatically – from a handful of defense systems against phage known until 2018 (most notably CRISPR and restriction-modification enzymes-RM) to 152 large families^1^. In parallel, our understanding of the mechanisms which phages use to circumvent this defense is quickly catching up and currently amounts to 156 known cases^3,4^. Although most of them are anti-CRISPR or anti-RM proteins^3,4^, a few dozen anti-defense proteins against other systems have recently been reported. Most directly counteract the defense system^5^, by blocking its activity^6,7^ or sequestering its product^8,9^, while others repair the damage caused by the defense system^10^. In all cases though, known anti-defenses counteract a single bacterial defense system, or more than one system when those use the same molecule^11^ or target the same host cellular machinery^10^.

Due to the short reproduction cycles of phages, defense and counter-defense mechanisms need to be fast^12^. Protein post-translational modifications (PTMs) are often used by cellular systems to rapidly sense and react to perturbations, and have been prevalently found in host-pathogen interfaces^13^. PTMs are also emerging as important in phage-bacterial interactions^14^ – for example acetylation deactivates host CRISPR defenses^15^, ubiquitin-like modification interferes with phage assembly^16^, and ADP-ribosylation has been long known to affect host transcription^17^ and more recently, translation^18^. Protein phosphorylation has also been demonstrated to be used as a bacterial host defense^19,20^ and very recently, also as phage counter-defense mechanism^21^.

Bacteriophage T7 infects *Escherichia coli* and encodes a serine-threonine (S/T) protein kinase known as gp0.7 or T7K (gene *0.7*)^22–24^. Its kinase activity was first described in the 1970s^24^, with subsequent studies identifying a small number of substrates, including the kinase itself^25^ and host proteins involved in transcription^26–28^, translation^29,30^, and nucleic acid processing^28,30,31^. The proposed model has been that T7K-mediated phosphorylation is directed to specific host proteins to hijack those cellular machineries for phage reproduction, by shutting off/modulating host transcription^26,27,32,33^ and stabilizing phage mRNAs^28^. However, a clear understanding of the function of T7K remains elusive. The kinase is dispensable in standard growth conditions with mild negative phenotypes in nutrient-poor media and heat stress^34^, and deactivates itself by phosphorylation early in the infection cycle^25^. Curiously, it also helps T7 to infect *E. coli* armed with plasmids encoding the channel-forming colicin-1b toxin^35–37^. Finally, although the C-terminal so-called “shut-off” domain of T7K is not required for kinase activity, it has been reported to be highly toxic and capable of inhibiting host transcription independently of the kinase domain^24,38,39^.

Here, we took advantage of modern phosphoproteomics to reinvestigate the activity and function of T7K. In contrast to the current model, we discovered that T7K acts as a hyper-promiscuous dual-specificity kinase, which targets the vast majority of phage and host proteins during a short burst of activity. Although hyper-promiscuous, T7K gains preference towards nucleic acid-binding proteins through its C-terminal domain, which we established binds to DNA. Preferential targets can be stoichiometrically phosphorylated, and may hence lose activity. Since many defense systems sense and/or target nucleic acids, we assessed T7K’s role upon infection of *E. coli* armed with different defense systems with DNA/RNA-binding activities. We showed that T7K-mediated phosphorylation at specific sites of host defense systems, such as Retron-Eco9^40^ and DarTG1^41^, abolishes their ability to defend against T7 infection. Finally, by assessing the role of T7K during infection of a library of natural *E. coli* isolates, we deduced that T7K has a broad-range impact on host defense mechanisms constituting a truly general anti-defense mechanism of phages.

## Results

### The T7 kinase hyper-phosphorylates the *E. coli* proteome during early infection stages

When applying a bacterial-adapted phosphoproteomics approach^42^ to study phage infection, we discovered striking hyper-phosphorylation of both host and phage proteomes during T7 infection. We detected 12,135 phosphopeptides in the wild-type (WT) T7 infection, compared to a few hundred in the uninfected and T7K knockout mutant controls (T7Δ*0.7::cmk,* hereafter abbreviated as Δ*0.7*)^43^ **(Fig. 1a-b, Supplementary Table 1)**. This implied that the phosphorylation was driven by T7K. The number of phosphorylation sites detected during T7 infection exceeds by several-fold the number of sites identified in any condition probed in *E. coli* before, and is ∼ 3-fold more than the total number of reported sites in *E. coli*^42,44–46^ **(Extended Data Fig. 1a-b, Supplementary Table 1)**. Since the majority of phosphorylation events occurred on sites and proteins that had not been previously reported, we reasoned that the T7K itself deposits these modifications. Phosphorylation sites included not only the expected serine (S) and threonine (T) residues^47^, but also tyrosine (Y) residues **(Fig. 1c)**, suggesting that T7K is a dual-specificity, highly promiscuous kinase. In agreement with this, FoldSeek^48^ predicted that the T7K domain’s tertiary structure resembled many dual-specificity kinases such as human CLK1, CLK2, DYRK1A **(Supplementary Table 2)**, and the top PFAM hit to the kinase’s orthologous protein group (PHROG^49^, phrog_2828; phylogeny of other member proteins shown in **Extended Data Fig. 1c**) was the dual-specificity protein kinase domain PF07714.16^49^ **(Supplementary Table 2).** Notably, no *close* homologues of T7K were found outside of phages **(Extended Data Fig. 1c, Supplementary Table 2).**

**Figure 1:**
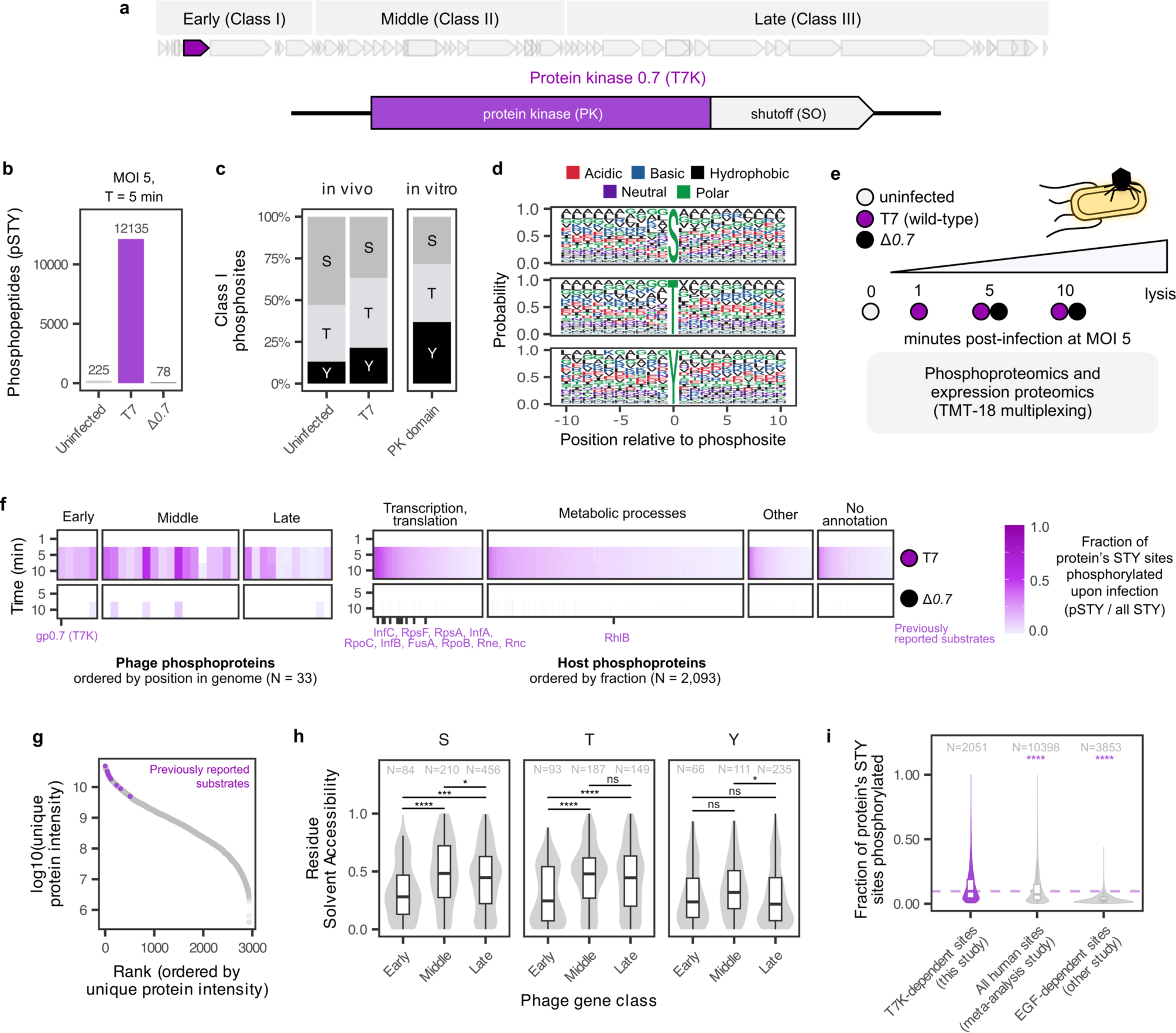
T7K hyper-phosphorylates the host and phage proteomes on serine, threonine and tyrosine residues. **(a)** Phage T7 genome with gene expression classes^52^. Gene *0.7* locus, encoding the protein kinase “T7K”, is highlighted and T7K protein domain architecture is depicted. **(b)** Infection by wild-type but not Δ*0.7* causes a massive phosphorylation of the *E. coli* proteome. Total number of unique serine (S), threonine (T) and tyrosine (Y) phosphopeptides identified from 3 biological replicates using phosphoenrichment and label-free MS/MS analysis. MOI = multiplicity of infection. **(c)** T7K is a dual specificity kinase. The proportion of class I phosphosites (PTMProphet localization score ≥ 0.75) localized to S, T and Y residues, identified in uninfected or T7 infected *E. coli* (panel **b**), “in vivo”, or from an in vitro phosphorylation assay using purified recombinant T7K kinase domain (6xHis-TEV-PK) applied to a tryptic digest of *E. coli* lysate. **(d)** T7K shows no sequence motif preference. Sequence motif logo plot centered around class I phosphorylated S, T or Y sites from the in vitro phosphorylation assay, using a 10-mer window of the surrounding residues. **(e)** Experimental design for the time-resolved T7 and Δ*0.7 E. coli* infection experiment with matched (phospho)proteome data (3 biological replicates) presented in (f). **(f)** T7K is responsible for rapid phosphorylation of the majority of the host and phage proteome. Fraction of significantly (limma, >2-fold, adjusted p-value<0.05; compared to uninfected cells) phosphorylated S/T/Y residues among all possible S/T/Y calculated per protein across the phage and host proteome during the time course infection. Proteins with at least one phosphosite are shown. Ticks and protein names indicate the previously reported phage and host substrates. Phage and host proteins are grouped according to their gene expression classes and GO slim biological process categories, respectively. **(g)** Previously reported T7K substates (purple) are among the most abundant *E. coli* proteins, as shown by ranking protein intensities from all time-course proteome measurements. **(h)** Accessible phosphorylatable residues are depleted from the surface of phage early and middle proteins. The solvent accessibility (RSA) values of S, T or Y residues within phage T7 proteins, estimated using AlphaFold3 structures. 1 means completely surface exposed and 0 means completely buried. Asterisks indicate statistical significance (unpaired t-test); * – p-value≤0.05, *** – p-value≤0.001, **** – p-value≤0.0001, ns – not significant, p-value≥0.05. **(i)** T7K targets a significantly larger faction of the proteome than all human kinases together. The fraction of phosphorylated S, T and Y residues per phosphoprotein compared between different datasets. “T7K-dependent sites” (purple) refers to phosphosites of host proteins which were significantly upregulated (limma, >2-fold, adjusted p-value<0.05) in wild-type T7 infection compared to uninfected and to Δ*0.7* at any matched time-point. “All human sites” refers to all phosphosites reported in a recent meta-analysis of the human phosphoproteome^85^. “EGF-dependent sites” refers to sites from a recent study on phosphoregulation in human HEK293F cells after EGF stimulation^86^. Comparisons made using unpaired two-sample Wilcoxon signed-rank tests (asterisks/significance as in **h**). N indicates number of phosphoproteins.

To validate the dual-specificity and high promiscuity of T7K, we performed in vitro phosphorylation of tryptic digests of *E. coli* proteins with the purified and dephosphorylated T7K kinase domain. This led to the identification of 12,642 phosphopeptides with phosphosites being evenly distributed across S, T and Y residues **(Fig. 1c)**. The proportion of tyrosine phosphorylation was higher in vitro compared to in vivo T7 infection **(Fig. 1c)**, likely reflecting the fact that Y residues are often buried in folded proteins^50^. Importantly, T7K exhibited no specificity whatsoever in the S, T or Y motifs it recognized **(Fig. 1d, Extended Data Fig. 2)**, which is consistent with high kinase promiscuity.

To better understand the dynamics of T7K, we monitored in parallel phosphopeptide and protein abundance in *E. coli* K-12 during WT and Δ*0.7* T7 phage infections. Since the T7 infection cycle is rapid (∼17 min to cell lysis), and the kinase is known to autophosphorylate within 6 minutes after infection and lose activity^25,47^, we collected samples just before infection (time-point 0) and 1, 5, or 10 minutes post-infection **(Fig. 1e-f, Extended Data Fig. 3)**. Although we did not identify any upregulated phosphopeptides after 1 minute of infection, consistent with T7K and overall phage proteins being barely expressed at this time-point **(Extended Data Fig. 4)**, we identified a massive phosphorylation increase in the later time-points, with most having occurred by 5 minutes **(Extended Data Fig. 3a-b)**. Collectively we quantified 15,697 phosphopeptides. Of these, 15,399 could be uniquely mapped to individual proteins, representing 11,057 phosphosites on 2,126 phosphoproteins (2,093 from host and 33 from phage) **(Supplementary Table 3)**. 14,663 unique phosphopeptides were T7K-dependent, i.e. significantly upregulated compared to uninfected and to Δ*0.7*, at any matched time-point **(Extended Data Fig. 3b)**, encompassing the vast majority of both host (2,051 out of 2,930 proteins, 70%) and phage (33 out of 48 proteins, 69%) expressed proteins **(Supplementary Table 3)**. Many host proteins were phosphorylated to high density (fraction of detectable S, T and Y residues that were phosphorylated), showcasing the processivity and promiscuity of T7K. This includes all 11 previously reported T7K-substrates^25–30^, which were amongst the most abundant proteins in the cell **(Fig. 1g).** As expected from the cytoplasmic action of the T7K, cell envelope and membrane proteins were largely depleted of phosphorylation **(Supplementary Table 3)**. In contrast to pervasive host protein phosphorylation, only 10 host proteins significantly changed in abundance in the absence of the kinase (comparing T7 and Δ*0.7* at 5 or 10 min, limma, adjusted p-value < 0.05, fold-change <0.5 or >1.5) **(Extended Data Fig. 4a-b)**. This included down-regulation of members of the colanic acid biosynthesis pathway, which protects against phage infection^51^ **(Extended Data Fig. 4c)**.

In terms of phage proteins, late (Class III)^52^ T7 proteins had lower phosphorylation densities than early or middle ones **(Fig. 1f)**, aligning with the previously described temporal regulation of kinase autoinhibition^25,47^. Interestingly, although early (Class I) T7 proteins^52^ were heavily phosphorylated, their sequences were selected to contain significantly less surface-accessible S and T residues **(Fig. 1h)**. This underlines the strong evolutionary pressure that the hyper-promiscuity of T7K puts on the T7 genome. We observed no such depletion of Y surface-accessible residues in early T7 proteins, suggesting they are less important for protein function and not subject to purifying selection.

Only a very small number of unique phosphopeptides (N = 15) exhibited a significant increase from 5 to 10 minutes post-infection **(Extended Data Fig. 3c)**, in agreement with the kinase being shut down by autophosphorylation at this point. In addition to the previously reported site S227^25^, we quantified phosphosites on T7K (Y122, Y150 and T346); and although the inhibitory S216^25^ was identified as a phosphosite in both label-free experiments **(Figure 1b,c)**, we could not confidently quantify it in the time-course **(Supplementary Table 3)**. In contrast to protein phosphorylation, protein abundance of middle and late expressed phage proteins increased between 5 and 10 minutes after T7 infection **(Extended Data Fig. 4d, Supplementary Table 3)**, confirming that phage infection is still actively progressing between these two time-points. Consistent with previous reports that a lack of the kinase leads to slightly slower T7 infection kinetics^34^, we observed lower middle and late protein abundances for Δ*0.7* infection, especially at the 5-minute time-point **(Extended Data Fig. 4d)**.

Taken together, we found an unprecedented extent of phosphorylation across the host and the phage (early and middle) proteomes within the first roughly 5 mins of infection, driven by the dual-specificity T7K. The majority of phosphorylation sites have never been observed before in *E. coli*. This degree of phosphorylation and processivity by one kinase exceeds what has been collectively reported in human cells over the years for all their encoded kinases (more than 500)^53^ **(Fig. 1i, Extended Data Fig. 1d)**. In contrast to this vast phosphoproteome remodeling, the *E. coli* proteome remains largely unaffected by T7K.

### The T7 kinase gains specificity to nucleic acid-binding substrates via its C-terminal domain

Phosphorylation can be used for signaling and to trigger downstream effects in the fate of proteins (e.g. change of localization or stability^54^), but if it is to directly change protein activity, then it needs to reach close to stoichiometric levels. Although T7K caused massive host proteome phosphorylation and appeared hyper-promiscuous in vivo and in vitro, we wondered whether any of these sites reached stoichiometric levels. However, measuring the fraction of a protein’s population that is phosphorylated is difficult, as phosphopeptides cannot be directly compared to their non-modified counterparts due to different peptide ionization efficiencies. To overcome this limitation, we compared the intensities of unmodified peptides before and after treatment of the peptide sample with a phosphatase enzyme, whereby the relative increase of the unmodified peptide intensity reflects the proportion of the peptide that was initially phosphorylated **(Fig. 2a).** When comparing unmodified peptide intensities in T7 and Δ*0.7* infected *E. coli* samples (10 minutes post-infection), we could determine robust Phosphatase/Control ratios (see Methods) for 4,355 peptides, of which 4,280 represented host proteins **(Extended Data Fig. 5a**, **Supplementary Table 4)**. We then identified peptides with significantly higher Phosphatase/Control ratios in the WT over the Δ*0.7* T7 infection (limma, adjusted p-value < 0.05 and fold-change > 1.5) **(Fig. 2b)**, which pointed to highly stoichiometric events **(Supplementary Table 4**). Peptides belonging to 82 host proteins, including 4/11 known substrates (bold), exhibited highly stoichiometric phosphorylation. Note that this analysis has false negatives, as robust ratios for lower abundance (phospho)peptides are harder to measure. Overall, there were many abundant proteins which were not stoichiometrically phosphorylated, including several previously identified substrates **(Extended Data Fig. 5b)**. Converting these ratios to actual stoichiometry values showed more clearly the highly stoichiometric phosphorylation events **(Fig. 2c)**. When examining whether the proteins corresponding to these peptides were enriched in specific protein classes using GO-term analysis, we found a significant enrichment in nucleic acid, DNA, rRNA, and RNA-binding molecular functions **(Fig. 2d, Extended Data Fig. 5c)**. While many of the previously reported T7K-substrates were rRNA and RNA binding proteins, the extensive number of DNA binding proteins with high phosphorylation stoichiometry was intriguing, as the only previously known substrates of this category were RNA polymerase subunits. Among the several transcription factors with stoichiometric phosphorylation was RcsB, which is known to activate colanic acid biosynthesis and hence influence biofilm production^51,55^. This was consistent with the proteomics results for changes in host protein abundance **(Extended Data Fig. 4c)**, and with T7K silencing this well-known bacterial defense against phage infection^55–58^.

**Figure 2:**
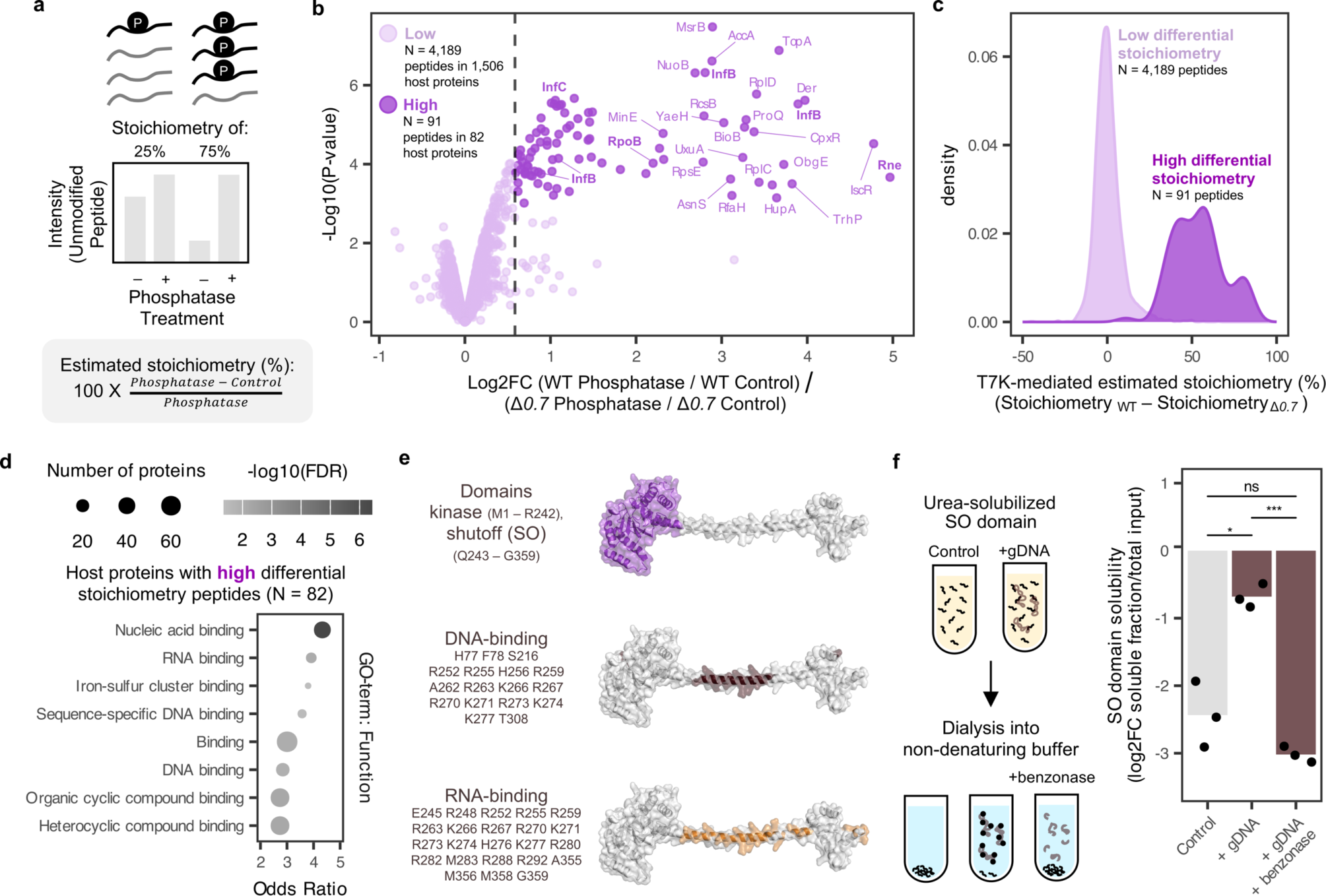
T7K hyperactivity is channeled towards nucleic acid-binding proteins due to direct interaction with DNA. **(a)** Schematic of how stoichiometry of phosphorylation sites can be experimentally estimated. **(b-c)** Quantitative analysis of phosphorylation in wild-type vs Δ*0.7* infected cells (T=10 minutes of infection, MOI=5, N=3 biological replicates) reveals high stoichiometry for specific host proteins – shown as volcano plots (**b**) and distributions of estimated stoichiometries (**c**). Only peptides uniquely mapping to host proteins, encompassing previously identified phosphosites from the time-course analysis (Fig. 1e), and with quality Phosphatase/Control ratios (see Methods), were assessed. Dark and light purple indicate peptides with a “High” (limma adjusted p-values<0.05, fold-change (FC)>1.5) and “Low” differential stoichiometry, respectively. The corresponding proteins are shown for peptides with high differential stoichiometry in **b**; bold font denotes previously described substrates. **(d)** Stoichiometrically phosphorylated substrates are enriched for nucleic-acid binding functions. GO-term analysis results for high stoichiometry proteins in **b** – showing only significant Function terms in which at least one category had an odds ratio >1 and number of genes >10. All identified proteins with unique peptides were used as background. **(e)** AlphaFold3^59^ structural prediction of the full-length T7K protein. A long positively charged alpha helix within the T7K shutoff (SO) domain is predicted to bind DNA and RNA by GraphBind^60^. **(f)** The T7K shutoff (SO) domain binds DNA. Recombinant 6xHis-TEV-SO (SO) is insoluble (Extended Data Fig. 6b). Urea-solubilized recombinant SO folded into a soluble protein in non-denaturing buffer in the presence, but not in the absence of genomic DNA (gDNA). Benzonase treatment reverted solubility, indicating that the SO domain assumes stable confirmation only when bound to DNA. Relative SO-domain solubility (abundance in the soluble fraction compared to the total input) was measured using TMT-multiplexed MS and shown as bar plot (N = 3 independent biological replicates; unpaired t-test analysis – ns (non-significant), * – p-value≤0.05, *** – p-value≤0.001).

Given the high number of stoichiometrically phosphorylated nucleic acid binding substrates (42/82 proteins), we reasoned that T7K may bind to nucleic acids and thereby localize its otherwise non-specific activity to these proteins. We generated an AlphaFold3^59^ model for the full-length T7K protein **(Extended Data Fig. 6a)** and analyzed it using the GraphBind tool^60^, which predicts the likelihood and position of RNA and DNA binding sites within protein structures. This revealed that the C-terminal domain (residues Q243 - G359) contained an extended alpha-helix highly enriched in basic residues predicted to bind both RNA and DNA **(Fig. 2e).** The C-terminal domain is not essential for the kinase activity^25,33^, but is known to be highly toxic and to be able to “shut-off” host transcription without the kinase domain^61^. Transcriptional shut-off by the C-terminus of T7K has been previously proposed to be mediated via tight DNA binding^33^, similarly to eukaryotic protamines.

In order to test T7K’s ability to bind DNA, we sought to purify the protein. We failed to recombinantly express full-length T7K (even with a tight inducible expression system), in line with previous studies reporting its toxicity^25^, but were able to individually express the protein kinase (PK) domain (residues 1-242) and C-terminal shutoff (SO) domain (residues 243-359). Although we were able to purify the PK domain, we were unable to obtain the SO domain in soluble form **(Extended Data Fig. 6b)**. We hypothesized that its folding may require direct interaction with nucleic acids, perhaps in a disorder-to-order transition, and that DNase I treatment during protein purification was causing the protein to precipitate. To test our hypothesis, we isolated the recombinant T7K SO domain (6xHis-TEV-SO) from insoluble inclusion bodies, and performed in vitro refolding. In this case, the urea-solubilized (denatured) protein was refolded by slow dialysis and buffer-exchange to native conditions (Methods) in the presence or absence of isolated *E. coli* genomic DNA (gDNA) **(Fig. 2f**). We then used quantitative mass spectrometry to measure its abundance in the isolated soluble fraction, and compared it to the total input. We found that the presence of gDNA during 6xHis-TEV-SO refolding significantly increased protein solubility. Furthermore, treating the gDNA sample after dialysis with benzonase, a nuclease that can degrade DNA, reversed this effect **(Fig. 2f)**. These results indicate that DNA plays a critical role in the stability of the SO domain, presumably via direct interactions between its positively-charged alpha-helix and DNA.

### The T7 kinase weakens the activity of phage defense systems by phosphorylation

Although T7K has a very broad specificity, it can stoichiometrically phosphorylate nucleic acid-binding proteins. As such proteins would likely be inactivated, we wondered why T7 would do this, as it still depends on intact host functions (e.g., translation) to complete its infection cycle. Prompted by the observation that T7K-mediated phosphorylation muted the Rcs response and colanic acid production **(Fig. 2b**, **Extended Data Fig. 4c)**, a broad-spectrum defense for phage infection^55–58^, we hypothesized that the T7K specificity to DNA-or RNA-binding proteins may be acting as a general deactivating mechanism for bacterial defense systems against phage. A plethora of such systems exists in bacteria, with most known ones sensing or targeting DNA, RNA, and/or their bound proteins^2,12,62,63^. Since such defense systems are scarce in *E. coli* K-12^64^, a historical host for phage biology, and other lab *E. coli* strains which have been chosen for their ability to take up foreign DNA, this would also explain the lack of phenotypes for T7K in the past.

To probe further whether T7K has an insofar undescribed functional role in the interaction with phage defense systems, we selected 12 systems with different DNA/RNA-links and one without **(Supplementary Table 5)**. Seven were retrons, which contain multicopy single-stranded DNA (msDNA) or DNA-RNA hybrids as a structural part of an antitoxin that keeps inactive cognate toxic effectors that defend against phage infections though abortive infection^40,65–67^. We expressed these defense systems under their native or arabinose-inducible (pBAD) promoters in *E. coli* K-12 (BW25113), and infected with T7 WT or Δ*0.7* at three different multiplicities of infection (MOIs: 5, 10^-2^, 10^-5^), while monitoring bacterial growth **(Fig. 3a, Extended Data Fig. 7)**. T7 infection led to reproducible culture collapse within one hour of infection, even for the lowest MOI (10^-5^), due to the rapid infection cycle (17 min) and a burst size of ∼180 virions per cycle^68^. We found three systems (retrons Eco6 and Eco9, and DarTG1) defending against T7 phage infection, the first two in line with previous reports^40,66^. Two out of three (Retron-Eco9 and DarTG1) seemed to be counteracted by T7K **(Fig. 3b).** All other systems did not defend at all, and deleting T7K did not alter this outcome **(Extended Data Fig. 7a-b)**. To gain better resolution for the T7K counter-defense effect, we tested the three defending systems at a higher MOI range. Whereas Retron-Eco6 showed a negligible difference between T7 WT and Δ*0.7* infection **(Extended Data Fig. 7c)**, the presence of the kinase had strong effects for both Retron-Eco9 and DarTG1 **(Fig. 3b, Extended Data Fig. 7d-e)**.

**Figure 3:**
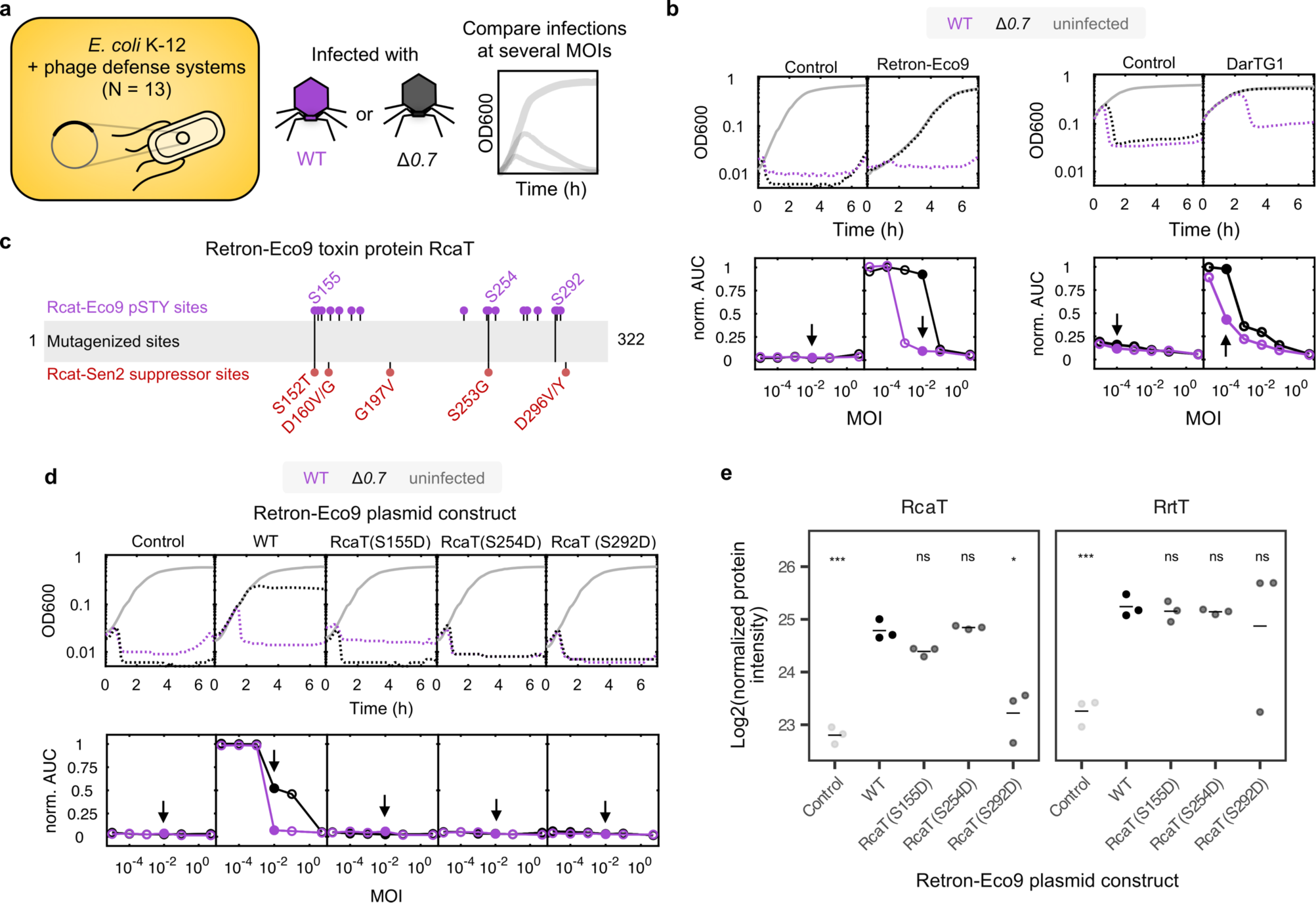
T7K weakens the activity of two DNA-related defense systems, Retron-Eco9 and DarTG, by phosphorylating them. **(a)** A panel of 13 bacterial defense systems was screened to assess their sensitivity to T7K. *E. coli* K-12 were transformed with each corresponding plasmid and infections at multiple multiplicities of infection (MOIs) with wild-type or the *0.7* (encoding T7K) knockout mutant were compared by monitoring bacterial growth and population lysis over >6 h (see Extended Data Fig. 7a-b for full results). **(b)** The defense systems Retron-Eco9 and DarTG1 defend against T7 and are sensitive to the T7K. For each system, a representative growth curve at a single MOI and normalized area under the curve (norm. AUC), summarizing growth curves across all MOIs, are shown. The MOI represented in the growth curve is indicated with an arrow in the corresponding norm. AUC plot. Control refers to *E. coli* K-12. N = 2 to 6 biological replicates (see Extended Data Fig. 7d-e for other replicates). **(c)** The RcaT encoded within the Retron-Eco9 defense system is heavily phosphorylated during T7 infection. Phosphoproteomics was performed at T = 5 mins after infection (N = 2 biological replicates). The positions of class I phosphosites (N = 17, unique, PTMProphet localization ≥ 0.75) are indicated along the length of full length RcaT as purple circles, and sites tested as phosphomimetic mutants (**d**) are named. The red dots indicate aligned residues in the homologous protein Retron-Sen2 RcaT that are critical for its function – as identified by a suppressor screen in^40^. **(d-e)** Single phosphomimetic mutants in Retron-Eco9 RcaT abolish its phage defense activity (**d**), without affecting protein stability in most cases (**e**). **(d)** Representative growth curves and norm. AUC plots for *E. coli* K-12 transformed with the various plasmid alleles presented as in panel **b** – N = 2 biological replicates (for other replicate see Extended Data Fig. 8a). **(e)** Protein stability of the mutant proteins was assessed by quantitative proteomics (N = 3 biological replicates). Unpaired t-tests comparing expression levels in each construct to the those in the WT Retron-Eco9 construct are shown, p-value depicted as in Fig. 2f.

To investigate whether the kinase phosphorylates and thereby inactivates these defense systems, we profiled their phosphorylation status during infection. Retron-Eco9 consists of a reverse transcriptase RrtT and its cognate msDNA, which together counteract the toxic RcaT effector^40^. Although the exact mechanism of RcaT toxicity is still unclear, structural insights into homologous retrons exist and possible cellular targets have been proposed^69,70^. When monitoring *E. coli* K-12 transformed with natively-expressed Retron-Eco9 after 5 minutes of infection with WT T7, we detected a large number of uniquely mapping, confidently-localized phosphosites within specific regions of the RcaT toxin protein (N = 17) **(Fig. 3b**, **Supplementary Table 6)**. In contrast, the RrtT protein had only four phosphorylated sites **(Supplementary Table 6)**. Two RcaT phosphosites, S155 and S254, aligned precisely with residues found to be critical for the function of the homologous RcaT protein (RcaT-Sen2) in a related retron, Retron-Sen2^40^ **(Fig. 3b).** We decided to construct phosphomimetic mutants (S→D; mimicking a constitutively phosphorylated state) of these RcaT sites (S155, S254), as well as of S181 and S292, which were aligned in close proximity to other functional residues in RcaT-Sen2. Although we repeatedly failed to produce the S181D mutated form, we were able to construct the three other alleles. All completely abrogated defense activity of Retron-Eco9 against T7 **(Fig. 3d, Extended Data Fig. 8a)**. Alleles S155D and S254D did this without largely affecting protein folding and abundance, whereas S292D presumably led to protein destabilization **(Fig. 3e)**. This implies that during phosphorylation of any of these residues, Retron-Eco9 RcaT loses its activity and/or stability, and hence the defense system is deactivated. The presence of several detrimental phosphorylation sites likely makes Retron-Eco9 particularly sensitive to the T7K anti-defense action.

DarTG1 is a prevalent toxin–antitoxin system, with its toxin DarT ADP-ribosylating thymidines and guanosines on phage DNA, and thereby inhibiting phage replication^41^. We took the same approach of phosphoproteomics followed by phosphomimetic mutants to probe the role of T7K on DarTG1. Both DarT and DarG were heavily phosphorylated upon T7 infection, with 19 and 18 phosphosites identified on each protein, respectively **(Supplementary Table 6)**. We chose to characterize the sites of the toxin DarT, and constructed three phosphomimetic mutants: T103D due to its structural proximity to the E152 active site for ADP-ribosylation, which is essential for the toxin’s activity^41^, and S65D and T167D, since they were close to highly conserved protein regions^41^. All three phosphomimetic DarT mutants were individually able to abolish DarTG1’s defense **(Extended Data Fig. 8b)**. However, in all three cases, the phosphomimetic mutations affected DarT, but not DarG, protein expression levels **(Extended Data Fig. 8c)**, likely because they affected its folding, structure and/or stability. Although this precludes us from definitively assessing the effect of phosphorylation on DarT protein activity, it is plausible that phosphorylation of folded DarT at these sites during infection affects DarT stability and hence defense.

In summary, we show that T7K can phosphorylate DNA-binding bacterial defense systems at or close to functionally relevant sites, and thereby disable their defense.

### The T7 kinase is likely a broadly-acting phage counter-defense system

The vast majority of the > 150 phage defense systems have been identified in the last 7 years and tested for defense against phage from medium-copy plasmids (10-15 copies/cell) in heterologous hosts^5,67,71^. Since T7K needs to stoichiometrically phosphorylate proteins to fully deactivate them, we reasoned that its anti-defense role may become more apparent when defense systems are expressed from the chromosome at native/lower levels of expression. In addition, bacterial strains carry multiple defense systems in their chromosome^1^. In order to test the function of T7K in a more physiologically relevant setup, we chose to infect a library of 513 *E. coli* natural isolates, comprising lab, commensal and pathogenic strains^72^. When annotating known defense systems present in these isolates using DefenseFinder^1,73^ and PADLOC^74^, it became apparent that they carried on average 9.0 and 9.9 defense systems, respectively **(Extended Data Fig. 9a**, **Supplementary Table 7)**.

We infected all *E. coli* strains during exponential growth with WT and Δ*0.7* T7 at 4 different MOIs (5, 10^-2^, 10^-3^ and 10^-4^), keeping a “no infection” control for each strain. We classified three key features: (i) the ability to be infected – by checking the response to WT T7 at MOI 5; (ii) the ability to defend – by checking whether lysis was prevented at MOIs lower than 5; and (iii) the presence of T7K-sensitive defense – by directly comparing the response to the respective MOIs between WT T7 and Δ*0.7* **(Fig. 4a, Extended Data Fig. 9b)**. Out of 513 strains (Methods), 54 (10.5%) were susceptible to T7 infection **(Fig. 4a, Extended Data Fig. 10)**. This low rate of infection was expected, as T7 infects only “rough” *E. coli* strains, i.e. those lacking O-antigen in their LPS^75^. Note that we cannot exclude that some of the non-infected strains are indeed infected, but carry defense systems that provide full defense. Out of the 54 infectable strains, 22 (4.3% of all strains) showed some degree of defense against T7 infection **(Fig. 4a, Extended Data Fig. 10c)**. We carefully reviewed these 22 strains at a higher MOI resolution (5, 10^-1^, 10^-2^, 10^-3^, 10^-4^, 10^-5^), and further excluded 5 strains with inconsistent phenotypes across biological replicates – mostly strains that now exhibited low/no infection **(Fig. 4a, Extended Data Fig. 11a-b)**. From the 17 “validated” strains left (infected by T7, but with signs of defense against T7), we found that the presence of the kinase (WT T7) reproducibly weakened the defense of six strains, strengthened the defense of one strain, and had negligible effects in ten strains **(Fig. 4b, Extended Data Fig. 11a-c)**. The effect size of the T7K-mediated weakening of defense spanned three orders of magnitude with two genetically diverse strains (H10407, IAI41) exhibiting effects of >1000-fold **(Fig. 4b)**, pointing to different defense systems being counteracted by T7K in these strains. Indeed, when looking into the defense systems harbored by these strains, there was no single common system that they encoded, which was absent from the ten no-effect/neutral strains **(Fig. 4c, Supplementary Table 7)**. Importantly, hit strains did also not contain any of the T7K-sensitive phage defense systems we discovered here (Retron-Eco9, DarTG), others functionally implicated with T7K (Retron-Eco2^76^), or recently described to be counteracted by the JSS1 phage protein kinase, which has some homology to T7K (Dnd, QatABCD, SIR2+HerA, DUF4297+HerA, CRISPR type I-E)^21^. This suggests that T7K inactivates a broad swath of diverse defense systems, which remain to be discovered in the future. It also implies that a defense system can evolve to evade T7K recognition and counter-defense (e.g. Dnd is present in the neutral strain, IAI55; CRISPR type I-E is present in most strains) or to sense T7K as a trigger (explaining why IAI15 can defend better against WT T7 than Δ*0.7* T7).

**Figure 4:**
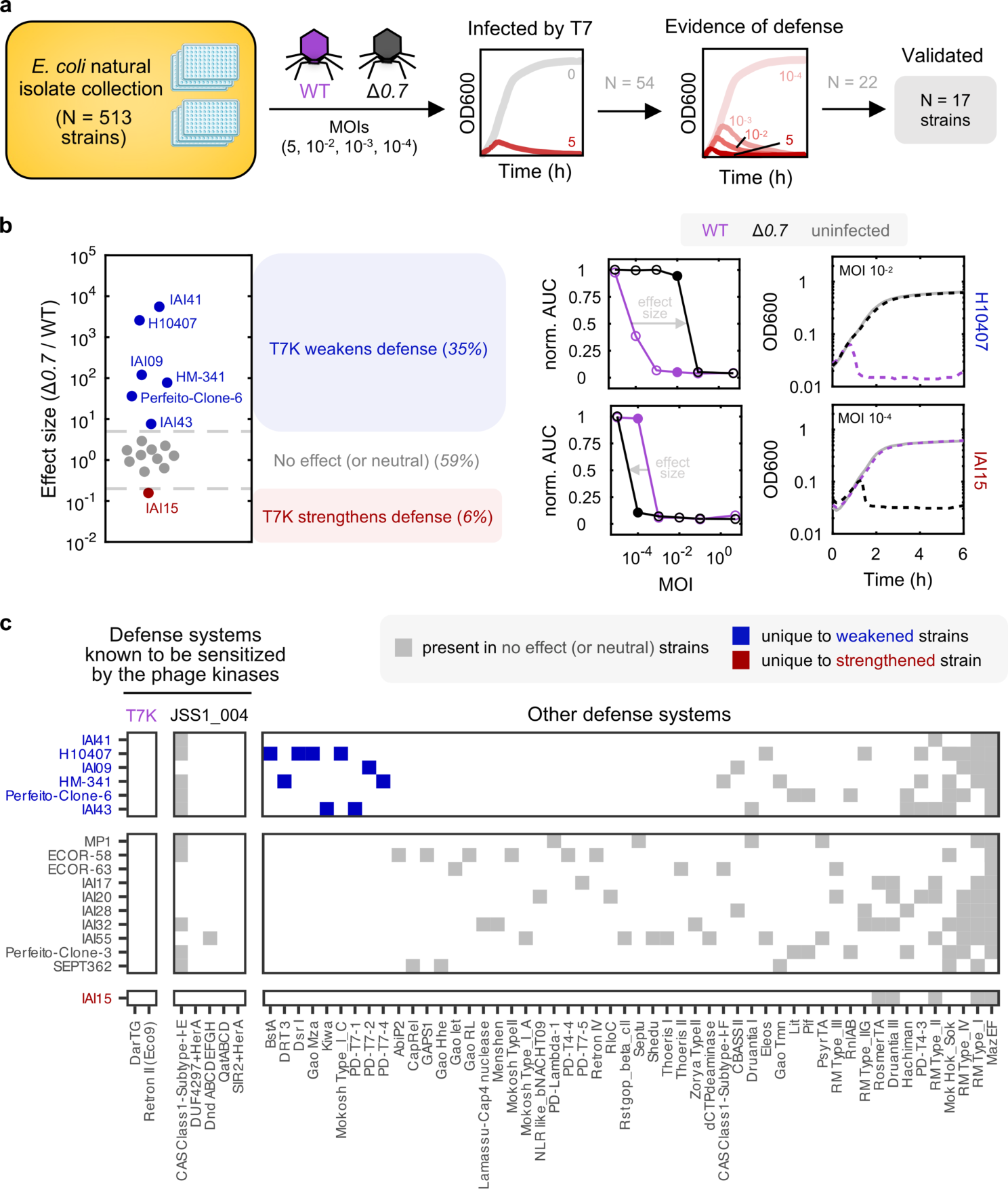
T7K helps T7 to infect diverse natural *E. coli* isolates, pointing to a broad counter-defense role. **(a)** Screen design to test for dependency on T7K during liquid infection assays of the ECOREF *E. coli* natural isolate library^72^ with WT and Δ*0.7* T7. Infectivity and evidence of defense were assessed in the screen (one replicate), and strains that passed were further validated using an extensive MOI and at least 2 biological replicates. **(b)** A considerable fraction of infectable strains by T7 exhibit defense that is sensitive to T7K. Effect size of sensitivity to T7K during infection for those strains that were infectable and had their evidence of defense was validated by independent assays. Effect sizes were estimated as the ratio (Δ*0.7*/WT) of the interpolated MOI at 50% AUC (see Methods). Strains were categorized as harboring defense systems that were weakened by T7K (blue, effect size ≥ 5), strengthened by T7K (red, effect size ≤ 0.2), or not affected (grey). An example growth curve at a single MOI and corresponding full norm. AUC graph are shown for a case of weakened and strengthened T7 defense. **(c)** T7K-sensitive natural isolates carry different defense systems, which are potential targets of T7K. DefenseFinder^1^ defense system profiles of validated strains grouped by their T7K-sensitivity categorization as defined in **b**. Defense systems are grouped by whether (i) we identified them to be T7K-dependent, (ii) they are sensitive to a homologous phage protein kinase JSS1_004^21^ or (iii) otherwise. The color indicates whether the system was uniquely found in weakened or strengthened strains, or whether they were also found in the no effect strains.

In summary, the results of the natural isolate screen **(Fig. 4)**, together with our targeted experiments **(Fig. 3)** and that of others from homologous phage kinases^21^, point to a broad counter-defense role for T7K against a plethora of bacterial immunity systems. This general counter-defense is achieved by targeting the highly promiscuous kinase activity of T7K via its C-terminal nucleic-acid binding domain towards DNA/RNA binding proteins, which are common in bacterial immunity systems.

## Discussion

In this work, we report that T7K acts as a hyper-promiscuous, dual-specificity kinase, which can phosphorylate any protein inside the infected cell. The substrate range of T7K exceeds that of any kinase studied before, and even that of all human kinases put together, with no motif whatsoever next to the phosphorylated S/T/Y residues. This loose cannon kinase gains specificity towards nucleic acid proteins through its C-terminal shutoff domain, and manages to stoichiometrically phosphorylate DNA/RNA-binding proteins within its short window of activity. The phage uses this stoichiometric phosphorylation to deactivate proteins sensing or attacking its DNA/RNA. To mitigate the dangers of such uncontrolled phosphorylation, T7 inactivates the kinase within 6 mins after infection via autophosphorylation^25,47^, and has depleted S/T exposed residues from the surface of its early and middle proteins. Such kinase activity has not been reported to our knowledge before, and sets a new paradigm of how phosphorylation can be used in living organisms – to deactivate molecular machines. Strikingly, and despite its vast impact on the cellular proteome, the role of the T7K has remained elusive for almost 50 years. This is because earlier tools lacked the sensitivity for identifying phosphorylated substrates^25–30^, but also due to the absence of strong phenotypes for the T7K mutant - having been tested in lab strains with poor phage defense repertoires.

Close homologues of T7K are only found in phages **(Supplementary Table 2)**. This suggests that the hyper-promiscuity of T7K phosphorylation could be an ancestral strategy that is incompatible with the selective S/T/Y signaling found in complex organisms. In the future, it would be interesting to study the activity of these close phage kinase homologues, and their relation to more distant dual-specificity kinases found in bacteria and eukaryotes, as this may give us insights into how such kinases may have become “domesticated” by tighter control of their activity and/or by gaining specificity. Other S/T kinases found in prophage regions of bacteria and known to be involved in phage defense such as Stk2^19^ have also been suggested to phosphorylate a broader substrate clientele than currently appreciated^20^. It remains to be further explored if such kinases are promiscuous, using sensitive phosphoproteomics methods like the one used in this study. More generally, S/T/Y phosphorylation has been considered to be less prevalent in prokaryote signaling (but with undoubted functional relevance^77^) based on the significantly fewer S/T or Y kinases and phosphosites found in bacteria compared to eukaryotes^77,78^. The present work shows that S/T/Y phosphorylation can be more pervasive in bacteria during phage infection than in eukaryotic cells.

We identified two defense systems that are sensitive to T7K, because they carry critical S/T residues on their surface. The decreased ability of the T7 phage lacking the kinase (Δ*0.7*) to bypass different natural *E. coli* isolates suggests that many more defense systems are counteracted by T7K. Recent studies corroborate our findings. When looking into T7 gene essentially using barcoded mutant libraries, Chen *et al.* identified an important role for T7K when infecting strains carrying Retron-Eco2^76^, presumably due to deactivation of the phage defense by T7K. Another study on *Salmonella* phage JSS1, which carries a kinase with 54% amino-acid sequence identity to T7K **(Extended Data Fig. 1c)**, demonstrated that this kinase could inactivate two other defense systems by phosphorylating them at critical residues (Dnd, QatABCD), and provided functional data for three further systems^21^. However, the extent of phosphorylation reported for this kinase was ∼20-fold lower than that of T7K, and defense systems were (some of) the strongest targets. It remains to be seen if the JSS1 kinase has lost some of the promiscuity seen for T7K and evolved specificity. Although exploring defense systems more systematically in the future will surely yield more T7K-sensitive systems, including possible ones that act on/bind to RNA, it is safe to say that this “sensitivity” will not be universal for all diverse members of a defense system class, as members may evolve to avoid critical STY residues on their surface. Some defense systems may have even adapted to use this (hyper)phosphorylation as a trigger, as indicated by one natural isolate that defended better when T7K was present.

Overall, this puts T7K and phosphorylation as one of the first general phage anti-defense mechanisms that can act independently of the catalytic activity of the defense system. The other known such general mechanism comes from Ocr, which is a protein encoded by a phage gene immediately next to the T7K (gp0.3) and blocks both R/M systems^79,80^ and BREX^81^ (as well as the host RNA polymerase^82^) via DNA mimicry^83^. Interestingly, other defense systems sense Ocr activity as a trigger^84^. Such general anti-defense mechanisms are bound to be more common, given the small size of phage genomes and the diversity of bacterial immune systems existing. Therefore, it is clear that PTM enzymes, with their prevalence in (pro)phages^14^ and their rapid responses (key for phage-bacterial interactions^12^), are good candidates for future investigations for general anti-defense mechanisms. Whether others are as hyper-promiscuous as T7K remains to be explored, and sensitive proteomics methods will be key for resolving these questions.

## Acknowledgments

We thank Udi Qimron and Ido Yosef for kindly providing the T7Δ*0.7::cmk* phage, Michael Laub and Michele LeRoux for the DarTG1 and CmdTAC plasmids, Inga Songailiene and Virginijus Šikšnys for the HEPN-MNT and ApeA plasmids, and Rotem Sorek for the Eco8 and Borvo plasmids. We thank A. Mateus for insightful discussions, and Jonathan Dworkin and Pedro Beltrao for reading and providing feedback for the manuscript. We would like also to thank all members of the Savitski and Typas groups for helpful discussions, Elisabeth Baland for the TacAT plasmid, the EMBL Proteomics Core Facility, in particular, Mandy Rettel and Jennifer Schwarz, and the EMBL Genomics Core Facility, in particular, Mireia Osuna Lopez, Hilal Ozgur and Vladimir Benes for expert help, and Elisabeth Baland for help with plasmid construction. This work was supported by the European Molecular Biology Laboratory. T.B. is supported by the EMBL EIPOD-LinC fellowship programme.

K.M. and C.M.P. were supported for part of the project by a fellowship from the EMBL Interdisciplinary Postdoc (EI3POD) programme under Marie Skłodowska-Curie Actions COFUND (MSCA COFUND 664726). M.M.S. is supported by the Allen Distinguished Investigator award through the Paul G. Allen Frontiers Group. M.G. was supported by the Deutsche Forschungsgemeinschaft (DFG, German Research Foundation) under Germany’s Excellence Strategy—EXC 2155—project number 390874280.

## Author contributions

A.T., C.M.P. and M.M.S conceived the study. T.B., K.M., C.M.P., A.T. and M.M.S. designed the study. A.T. and M.M.S. supervised the project. C.M.P., T.B., and I.B. performed and analyzed the proteomics experiments; K.M., T.B., A.L.J.Y., A.K., and J.B. performed and analyzed microbiology experiments; F.C. performed and analyzed the biochemical experiments. T.B. did computational analysis with contributions on proteomics data from M.L.B. and on genomics data from N.K. and M.G.. T.B., K.M., C.P., A.T., and M.M.S. wrote the manuscript with input from all authors. T.B. and K.M. designed and plotted the figures with inputs from C.M.P., A.T. and M.M.S. All authors approved the final version.

## Author information

Correspondence and requests for materials should be addressed to.

## Supplementary Information

**Supplementary Table 1** Phosphoproteome of phage infection (single time-point)

**Supplementary Table 2** Analyses of T7K structural and sequence homology

**Supplementary Table 3** Quantitative phosphoproteome and proteome analysis of phage infection (time-course)

**Supplementary Table 4** Estimation of phosphorylation stoichiometry

**Supplementary Table 5** Plasmid-based screen of T7K interaction with phage defense systems

**Supplementary Table 6** Retron-Eco9 and DarTG1 analyses

**Supplementary Table 7** Natural isolate library screen details and defense system characteristics

## Source Data

Source Main Fig. 1-4

Source Extended Data Fig. 1-11

Source Supplementary Table 1-7

## Methods

### Bacterial strains

All strains for the experiments were grown in Lysogeny Broth (LB) Lennox supplemented with 5 mM CaCl_2_ and 10 mM MgCl_2_, and the required antibiotics and incubated at 37°C with shaking. *Escherichia coli* K-12 (BW25113^91^) was used for most biological analyses, whilst the strains BL21-AI or NEB5ɑ were used for protein purification and cloning, respectively. The use of the ECOREF^72^ strain collection is described in detail in later sections. When culturing strains transformed with plasmids, the following antibiotic selection was used: 100 μg/ml spectinomycin, 25 µg/ml chloramphenicol, 100 μg/ml kanamycin.

### Phages

Phage T7 WT was acquired from DSMZ, T7Δ*0.7::cmk* (referred to as Δ*0.7* throughout the manuscript) was a gift from U. Qimron^43^. Our phage stocks were checked for the presence of the correct T7 genotype with the primers TAGCCAACACACTGAACGCT & CACCCGCAGATGACCTGTAA for T7 WT and TAGCCAACACACTGAACGCT & GCCAAAAGGCGCTCAAAGTT for T7Δ*0.7::cmk*. In addition, we confirmed that the known deletion mutant H1^92^ did not occur in our stocks, using the primers TAGCCAACACACTGAACGCT, CACCCGCAGATGACCTGTAA, and GCCAAAAGGCGCTCAAAGTT. Phages were propagated in the *E. coli* K-12 (BW25113) strain and cultured in LB, 5 mM CaCl_2_, 10 mM MgCl_2_, at 37°C, shaking. Lysates of overnight cultures infected with phage stocks were prepared by adding chloroform, vortexing, and centrifuging (4,000 xg, 4°C, 5 min) to remove debris. The lysate was transferred to fresh tubes, additional chloroform was added, and samples were vortexed and centrifuged again. Clarified lysates were stored at 4°C. Lysate titres were determined in duplicates using the double-agar overlay method^93^.

### Plasmids

For the study of recombinant T7K, the regions encompassing the N-terminal PK domain (residues 1-242) and the C-terminal SO domain of T7K (residues 243-359) were PCR-amplified from T7 genomic DNA using Q5 High-Fidelity DNA Polymerase (NEB), with primers designed to introduce a 6xHis-TEV N-terminal fusion and carrying overhangs to plasmid pET28a.

Almost all plasmids used in the T7K screen **(Supplementary Table 5)** are in the pTU175 backbone, which carries the low-copy number origin of replication pSC101, except for the Borvo and Retron-Eco8 defense systems, which were obtained from the lab of Rotem Sorek and kept in the pSG1-vector –expressed from their native promoter and with a p15a origin of replication^66,71^, and CmdTAC (PD-T4-9) defense system, which was obtained from the lab of Michael Laub, and was kept on its original plasmid pCD1–PD-T4-9 under its native promoter^94^. Plasmids containing Retrons-Eco1, - Eco9 and -Sen2 have been described before^40^. For Retrons-Eco3, -Eco6 and -Eco10, the retrons were amplified with their native promoters (500 bp upstream of the retron start) from strains from the ECOREF *E. coli* natural isolate library^72^ or from strain SC402 (Retron-Eco3)^40^, and cloned into pTU175. For the toxin–antitoxin system TacAT^95^, the protein-coding sequence was amplified from *Salmonella typhimurium* LT2 and cloned into the pTU175 backbone under the arabinose-inducible pBAD promoter. The HEPN-MNT^96^ and the ApeA^67^ defense systems were cloned into the pTU175 vector under the arabinose-inducible pBAD promoter (gift by Inga Songailiene - Virginijus Šikšnys lab). The DarTG1 defense system was cloned into the pTU175 vector under the arabinose-inducible pBAD promoter by Michele LeRoux^97^.

Phosphomimetic mutant constructs of RcaT (in Retron-Eco9) and DarT (in DarTG1) were generated using the Q5 Site-Directed Mutagenesis Kit (NEB), using the primers described in **Supplementary Table 6.** Plasmid edits were confirmed by Sanger sequencing and PacBio whole plasmid sequencing.

### Infection time-course sampling

To measure proteome and phosphoproteome changes in wild-type T7 and Δ*0.7* phage infections of *E. coli* K-12 (BW25113), a single colony was picked and grown overnight in LB, 10mM MgCl_2_, 5 mM CaCl_2_ and the following morning back diluted 1:1,000 into fresh LB, 10 mM MgCl_2_, 5 mM CaCl_2_ (300 ml). The bacteria were cultured at 37°C, shaking, with their optical density at 600 nm monitored over time until an OD600 of ∼0.5 (1cm pathlength). The culture was then split into six independent conical flasks and phage stock (either wild-type or Δ*0.7*, or no stock as the uninfected control) added to a final multiplicity of infection (MOI) of 5 and infections were allowed to proceed at 37°C, shaking. Uninfected samples were immediately placed on ice. To harvest at 1, 5, or 10 min post-infection for wild-type T7, or at 5 and 10 min for Δ*0.7*, the flasks were immediately transferred to a pre-cooled centrifuge and spun (4,000 x g, 5 min, 4°C). For each sample, the supernatant was removed and the cell pellet was resuspended in 1 ml ice-cold PBS, and then centrifuged once again (10,000 x g, 1 min, 4°C). The supernatant was aspirated and the pellet was flash-frozen in liquid nitrogen. In total, three biological replicates were performed (time-courses from three independent colonies prepared on different days).

### MS sample preparation (time-course)

The lysis buffer was composed of 4 M guanidinium isothiocyanate, 50 mM HEPES (2-[4-(2-hydroxyethyl)piperazin-1-yl]ethanesulfonic acid), 5 mM TCEP (tris(2-carboxyethyl)phosphine), 20 mM Chloroacetamide, 1% N-lauroylsarcosine, 5% isoamyl alcohol, and 40% acetonitrile, with the pH adjusted to 8.5 using 10 M NaOH. For cell lysis, a buffer volume of approximately five times the cell pellet volume was added. The samples were homogenised by pipetting, incubated on a shaker at room temperature for 1 h, and centrifuged at 16,000 x g for 10 min to remove cell debris and nucleic acid aggregates. Protein concentration was determined via tryptophan fluorescence, as previously described^98^. Samples were transferred to MultiScreenHTS-HV 96-well filter plates with 0.45 µm PVDF membranes (Merck Millipore), and ice-cold acetonitrile was added to each well to induce protein precipitation, achieving a final concentration of 80% acetonitrile. After a 10 min incubation, the samples were centrifuged, and the supernatant was discarded. The protein precipitates were then washed twice with 200 µl of 80% acetonitrile and twice with 200 µl of 70% ethanol, with each wash followed by centrifugation at 1,000 x g for 2 min.

Next, a digestion buffer containing 50 mM Triethylammonium bicarbonate buffer, and trypsin (TPCK-treated, Thermo Fisher Scientific) was added to the protein precipitates. The trypsin-to-protein ratio was set to 1:25 (w/w), with a maximum final protein concentration of 10 µg/µl. Tryptic digestion was performed overnight at room temperature with mild shaking (600 rpm). After digestion, the samples were dried using a vacuum concentrator.

For the parallel proteomics measurements, the full digest was TMT-labelled (see below) and then subjected to offline high pH fractionation (see below). For phosphoproteomics measurements phosphopeptide enrichment was performed (see below).

### Phosphopeptide enrichment

For phosphoproteomics studies (time-course, Retron-Eco9 and DarTG1 infections), phosphopeptide enrichment was performed. Lyophilised peptides were resuspended in a loading and washing buffer composed of 80% acetonitrile and 0.07% TFA, sonicated, and centrifuged at 16,000 x g. Phosphopeptide enrichment was carried out on the supernatant as described in ^99^ using the KingFisher Apex robot (Thermo Fisher Scientific) with 25 µl of Fe-NTA MagBeads (PureCube) per sample. After five washes with buffer A, phosphopeptides were eluted with 100 µl of 0.2% diethylamine in 50% acetonitrile, followed by lyophilisation.

The enriched phosphoproteomic time-course samples were TMT labelled (see below) and then subjected to offline PGC fractionation. The enriched Retron-Eco9 and DarTG1 infection samples were not TMT-multiplexed and were subjected to MS analysis without further peptide fractionation.

### MS sample preparation (Retron-Eco9 and DarTG1)

For phosphoproteomics studies, *E. coli* K-12 were transformed with plasmids encoding Retron-Eco9 (native expression) or DarTG1 (arabinose-inducible). Overnight cultures were started in LB, 10 mM MgCl_2_, 5 mM CaCl_2_, 100 μg/ml spectinomycin, and then from these a 50 ml overday day culture (1:2,000 back dilution) was started the following morning in fresh LB, 10 mM MgCl_2_, 5 mM CaCl_2_, 100 μg/ml spectinomycin. DarTG1 culture media had additional 0.2% (w/v) arabinose for the induction of DarTG1 expression. Cells were grown until OD600 of ∼0.05 (Retron-Eco9) or ∼0.6 (DarTG1) (pathlength = 1cm), infected with wild-type T7 at an MOI of 5, and then harvested after 5 min of infection, as described above in sections: MS sample preparation (time-course), Phosphopeptide enrichment. Two independent biological replicates were performed.

For experiments assessing phosphomutant protein stability (through abundance), cells transformed with either Retron-Eco9 or DarTG1 constructs (or their corresponding empty plasmids) were cultured as described above, but harvested (without phage infection) at OD600 ∼0.05 (Retron-Eco9) or ∼2 (DarTG1). Cells were harvested by centrifugation (5 min, 4,000 xg, 4°C), washed once with ice-cold PBS, centrifuged again and then resuspended in 5 pellet volumes of 2% (w/v) SDS. Samples were boiled for 5 min, and then subjected to 5 rounds of freeze-thaw before being processed for MS analysis (sample cleanup, proteolytic digestion, TMT-multiplexing, high pH fractionation - see below sections). Three independent biological replicates were performed.

### MS sample clean-up and proteolytic digestion

MS sample preparation and measurements were performed as in described ^100^. Protein digestion was performed using a modified SP3 protocol^101^. Samples were diluted to a final volume of 20 μl with 0.5% SDS and mixed with a paramagnetic bead slurry (10 μg beads (Sera-Mag Speed beads, Thermo Fischer Scientific) in 40 μl ethanol). The mixture was incubated at room temperature with shaking for 15 min. The beads now bound to the proteins were washed 4 times with 70% ethanol. Proteins on beads were reduced, alkylated and digested using 0.2 μg trypsin, 0.2 μg LysC, 1.7 mM TCEP and 5 mM chloroacetamide in 100 mM HEPES, pH 8. Following overnight incubation, the peptides were eluted from the beads, dried under vacuum, reconstituted in 10 μl of water and labelled with TMTpro, 18plex reagents and dissolved in acetonitrile at 1:15 (peptide:TMT weight ratio) for 1 h at room temperature. The labelling reaction was quenched with 4 μl of 5% hydroxylamine and the conditions were pooled together. The pooled sample was desalted with solid-phase extraction after acidification with 0.1% formic acid. The samples were loaded on a Waters OASIS HLB μelution plate (30 μm), washed twice with 0.05% formic acid and finally eluted in 100 μl of 80% acetonitrile containing 0.05% formic acid. The desalted peptides were dried under vacuum and dissolved for LC-MS analysis.

### Expression and purification of 6xHis-TEV-PK

Plasmid pET28a-6xHis-TEV-SO was transformed into *E. coli* BL21-AI. Cells were grown at 37°C in 1.2 l of ZYM-5052^102^, supplemented with 100 μg/ml kanamycin (Sigma-Aldrich) and 0.05% arabinose (Sigma-Aldrich), and cultures shifted to 30°C when OD600 reached ∼0.8-0.9 (pathlength = 1cm). Protein overproduction was carried out for further 18 h. Cells were harvested in a Beckman-Coulter JLA 8.1000 rotor (5,000 rpm, 30 min, 4°C) and pellets resuspended in 45 ml of lysis buffer (50 mM Tris pH 7.5, 5 mM EDTA) supplemented with EDTA-free protease inhibitor (Roche), 100 μg/ml DNase I (Sigma-Aldrich) and 0.5 mg/ml lysozyme (Sigma-Aldrich). Cells were lysed in a C3 Emulsiflex high-pressure homogenizer (Avestin) and insoluble material pelleted (80,000 x g, 20 min, 4°C). The pellet was resuspended in 30 ml of membrane solubilization buffer (50 mM Tris pH 7.5, 2% Triton X-100) and samples were pelleted again (80,000 x g, 20 min, 4°C). The pellet was then resuspended in high salt buffer (50 mM Tris pH 7.5, 1 M NaCl, 2% Triton X-100) and ultracentrifuged again (80,000 x g, 20 min, 4°C). Inclusion bodies were resuspended in high urea buffer (50 mM Tris pH 8.0, 7.6 M urea) and let solubilize at 4°C for 18 h with gentle shaking. Insoluble material was removed by ultracentrifugation (80,000 x g, 20 min, 4°C) and urea-solubilized protein was stored at −80°C.

### In vitro phosphorylation assay

To remove autoinhibitory phosphosites on T7K and enable the study of its activity in vitro, dephosphorylation of purified T7K kinase domain was performed. The reaction was performed in a total volume of 4.5 ml. Purified 6xHis-TEV-PK was supplemented with 10 mM MnCl_2_ and 4,000 U of lambda protein phosphatase (NEB), and incubated at 30°C. After 3 h, another 4,000 U of lambda protein phosphatase were added, and the sample was incubated at 30°C for another 3 h. To remove lambda protein phosphatase, the mixture was applied to 500 μl of pre-equilibrated Ni-NTA agarose beads (Qiagen) on a gravity flow column. Beads were washed with 20 ml of binding buffer (20 mM Tris pH 7.5, 500 mM NaCl, 5 mM imidazole), then 6xHis-TEV-PK eluted in 5 ml of elution buffer (20 mM Tris pH 7.5, 500 mM NaCl, 1 M imidazole), collecting 500 μl aliquots. Fractions with the highest purity and yield were pooled and dialyzed twice against 2 l of dialysis buffer (20 mM Tris pH 7.5, 500 mM NaCl, 1 mM DTT, 1 mM EDTA). The protein was stored at −80°C in 30% glycerol at a final concentration of 0.5 mg/ml.

For the in vitro phosphorylation assay, 500 μg of *E. coli* K-12 tryptic digest per replicate were resuspended in a buffer containing 50 mM HEPES pH 7.5, 100 mM NaCl, 5 mM Mg-ATP and 1 mM TCEP. Subsequently, 5 ug of the purified (and dephosphorylated) kinase domain was added, and the reaction was incubated at 37°C for 4 h. Then, the peptides were lyophilized before phosphopeptide enrichment and label-free LC-MS/MS. In total, three biological replicates were performed.

### Phosphatase experiment to estimate phosphorylation stoichiometry

Tryptic peptides (40 µg for each of the three replicates) corresponding to either *E. coli* infected with T7 WT or Δ*0.7* (at T = 10 min post-infection, three biological replicates) were subjected to alkaline phosphatase treatment (5 units of FastAP per sample, Thermo Fisher Scientific) in a volume of 50 µl of phosphatase buffer, at 37°C for 20 h. As a control, the same amount of tryptic peptides were incubated in the same conditions without phosphatase. After incubation, samples were lyophilized and desalted using t-C18 Sep-Pak columns (Waters) before TMT labeling and offline PGC fractionation.

### 6xHis-TEV-SO in vitro refolding assay

Genomic DNA (gDNA) was isolated from 50 ml of *E. coli* BW25113 overnight culture using DNeasy Blood & Tissue Kit (Qiagen) and concentrated to ∼0.8 mg/ml in water in a vacuum concentrator. Refolding reactions were assembled as follows: gDNA (80 μg) or an equal volume of water were mixed with 40 μg of urea-solubilized 6xHis-TEV-SO in 25 mM Tris pH 8.0, 3.8 M urea, in a total volume of 200 μl. Mixtures were transferred to 6-8 kDa molecular weight cutoff D-Tube Dialyzer Mini tubes (Merck-Millipore) and dialyzed for 18 h at 4°C with gentle stirring in refolding buffer (25 mM Bis-Tris pH 6.6, 500 mM NaCl, 1 mM MgCl_2_). Aliquots (15 μl) of total material (input fraction) were collected after dialysis, then the rest of the sample was centrifuged (16,000 x g, 45 min, 4°C) and supernatants (soluble fraction) were collected. Benzonase-treated controls were prepared as follows: 50 μl aliquots of soluble fraction from samples containing gDNA were supplemented with 125 U of benzonase (Merck-Millipore) and 10 mM MgCl_2_ and incubated at 37°C for 4 h, then insoluble material removed by centrifugation (16,000 x g, 45 min, 4°C) and supernatants collected. In total, samples across three biological replicates were collected.

MS sample preparation of all fractions (input, soluble, benzonase-treated controls), including TMT-multiplexing was prepared as described above (Sample clean-up and proteolytic digestion). As there was some evidence of remnant protease from the genomic extraction kit, for protein digestion, the ‘nonspecific’ setting was used as protease with an allowance of maximum 2 missed cleavages requiring a minimum peptide length of 7 amino acids.

### TMT Labeling

For the full proteome measurements, 10 µg of peptides per condition were labeled, while for the phosphoproteome, the totality of enriched phosphopeptides was used. Peptides were resuspended in 10 µl of 100 mM HEPES (pH 8.5), and 4 µl of TMTPro reagent at a concentration of 20 µg/µ in acetonitrile was added. The labelling reaction was allowed to proceed for 1 h at room temperature, and then was quenched by the addition of 5 µl of 5% hydroxylamine for 15 min. Labeled peptides belonging to the same experiment were subsequently pooled and lyophilized.

Before fractionation, the phosphopeptides were resuspended in 50 µl of 10% TFA and desalted using in-house C18 stage tips^103^ packed with 1 mg of ReproSil-Pur 120 C18-AQ 5 µm material (Dr. Maisch) above a C18 resin plug (AttractSPE disks bio - C18, Affinisep), while in the case of the full proteome samples, the peptides were desalted using a Sep-pak t-C18 column (Waters).

### PGC-LC offline peptide fractionation

The samples were resuspended in 18 µl of buffer A (0.05% TFA in MS-grade water supplemented with 2% acetonitrile), with an injection volume set to 16 µl. Peptide separation was conducted using a Hypercarb column (100 mm length, 1.0 mm inner diameter, 3 µm particle size, Thermo Fisher Scientific) at 50°C, with a flow rate of 75 µl/min, on an Ultimate 3000 Liquid Chromatography system (Thermo Fisher Scientific). After a 1-min postinjection delay, a linear gradient separation was applied, increasing from 13% buffer B (0.05% TFA in acetonitrile) to 42% buffer B over 47 and 95 min for the phosphoproteome and full proteome samples, respectively. This was followed by an increase to 80% buffer B within 5 min. The column was then washed with 80% buffer B for 5 min before re-equilibration with 100% buffer A for another 5 min. Fractions were collected from 4.5 to 52.5 min and from 4.5 to 100.5 min at 2-min intervals, yielding 24 fractions and 48 fractions for the phosphoproteome and full proteome samples, respectively. These were then pooled into 12 or 24 fractions by combining each fraction with its corresponding n + 12 and n + 24 fraction for the phosphoproteome and full proteome samples, respectively. After pooling, the fractions were dried using a vacuum concentrator prior to LC-MS/MS analysis.

### High-pH offline peptide fractionation

Samples were fractionated using a reversed-phase C18 system running under high pH conditions, as described previously in ^100^. Briefly, this consisted of an 85 min gradient (mobile phase A: 20 mM ammonium formate (pH 10) and mobile phase B: acetonitrile) at a 0.1 ml min^−1^ starting at 0% B, followed by a linear increase to 35% B from 2 min to 60 min, with a subsequent increase to 85% B from up to 62 min and holding this up to 68 min, which was followed by a linear decrease to 0% B up to 70 min, finishing with a hold at this level until the end of the run. Fractions were collected every two min from 12 min to 70 min, and fractions were concatenated to make a total of 6 fractions (Retron-Eco9, DarTG1 expression analysis) or 24 fractions (phage infection time-course expression analysis) by combining each fraction with its corresponding n + 6 and n + 24 fraction.

### LC-MS/MS

Prior to injection, all samples were resuspended in a loading buffer composed of 1% TFA, 50 mM citric acid, and 2% acetonitrile in MS-grade water. Liquid chromatography separation was performed on an UltiMate 3000 RSLCnano system (Thermo Fisher Scientific). Peptides were initially trapped on a cartridge (Precolumn: C18 PepMap 100, 5 μm, 300 μm i.d. × 5 mm, 100 Å) before being separated on an analytical column (Waters nanoEase HSS C18 T3, 75 μm × 25 cm, 1.8 μm, 100 Å). Solvent A was composed of 0.1% formic acid with 3% DMSO in LC–MS-grade water, while solvent B contained 0.1% formic acid with 3% DMSO in LC–MS-grade acetonitrile. Peptides were loaded onto the trapping cartridge at 30 μL/min with solvent A for 5 min, then eluted at a constant flow rate of 300 nl/min using a linear gradient of buffer B, followed by an increase to 40% buffer B, a wash at 80% buffer B for 4 min, and re-equilibration to initial conditions. The linear gradient corresponded to an increase from 5% to 25% B in the case of label-free peptides, and from 7% to 27% B in the case of TMT-labeled peptides.

The LC system was coupled to either a Fusion Lumos Tribrid, an Exploris 480 mass or a Q-Exactive Plus mass spectrometer (Thermo Fisher Scientific), operated in positive ion mode. The mass spectrometers were operated in data-dependent acquisition mode with a maximum duty cycle time of 3 sec or a top 20 method, selecting precursors with charge states 2–7 and a minimum intensity of 2 × 10⁵ for subsequent HCD fragmentation. Peptide isolation was performed using the quadrupole with 0.7 m/z or 1.4 m/z isolation windows in the case of TMT-labeled and label-free samples, respectively. MS/MS spectra were acquired in profile mode using the Orbitrap, with a maximum injection time of 100 ms and an AGC target of 1 × 10⁵ charges.

### LC-MS/MS data analysis

For all proteomic analyses, a combined protein sequence database was constructed consisting of the Swissprot *Escherichia coli* strain K-12 database (4,401 entries, UP000000625) supplemented with a common protein contaminants database (118 entries). For analyses involving phage infections, this was further supplemented with the Swissprot phage T7 proteome (UP000000840, 57 entries). For analyses involving *E. coli* transformed with either defense system-containing plasmids, the database was also supplemented with the proteins encoded on each plasmid - DarTG1 (5 entries) and Retron-Eco9 (5 entries). For the shut-off domain solubility analysis, the Swissprot *Escherichia coli* strain K-12 database was supplemented with contaminants and additionally supplemented with the sequence for the recombinant T7K SO-domain protein.

Raw files were converted to mzmL files using MSConvert from Proteowizard^104^, using peak picking and keeping the 1,000 most intense peaks per spectrum. Files were then searched using MSFragger v4.0^105^ in Fragpipe v21 against the combined *E. coli*, T7 and contaminants database (see above, 4,576 sequences). The default MSFragger phospho or TMT16-phospho workflow was used, with a few modifications: oxidation on methionine (maximum 2 occurrences), phosphorylation on S/T/Y (maximum 3 occurrences) and peptide n-terminal TMT16 labeling (maximum 1 occurrence) were set as variable modifications, with a total of up to 5 variable modifications allowed per peptide. Lysine TMT16 labeling and cysteine carbamidomethylation were set as fixed modifications. Percolator^106^ was used for PSM validation and PTMProphet^87^ was used to determine site localization. An FDR cutoff of 1% was used. For the TMT quantification, Philosopher^107^ was used to extract MS1 and TMT intensities.

### Data processing and statistical analysis - general

For the summarisation of information at the phosphosite level, the highest localisation score for each site across redundant peptide spectrum matches (PSMs) was considered. Class I phosphosites were those that were uniquely mapping (only one protein in the FASTA database) with a PTMProphet^87^ phosphorylation localisation probability ≥0.75. The summarisation of quantitative information at the (phospho-)peptide levels (for phosphoproteomics and stoichiometry experiments) was performed by summing the TMT intensities of all the PSMs assigned to that specific (phospho-)peptide.

For TMT analyses, the PSM tables produced by FragPipe were filtered to keep PSMs having a purity value >0.5 with full quantification (signal in all utilised TMT channels). For all experiments, contaminant proteins were filtered out. For phosphoproteomics experiments, all phospho-modified peptides were considered; for the stoichiometry experiment, only unique peptides were considered; for protein expression analysis, only proteins with at least one unique peptide were considered.

Differential expression analysis was performed using the summed TMT reporter ion intensities (summarised at the Modified Peptide level) for phosphoproteomics and stoichiometry experiments, or using the protein intensities from MSFragger output for other protein expression experiments. Vsn-normalisation^89^ and differential expression analysis were used as implemented in the limma package (version 3.58.1)^108^ in R (version 4.3.3).

For the time-course experiments, vsn-normalisation was performed grouped by sample type (phage genotype + time-point) for phosphoproteomics, or for proteomics grouped by the protein’s species origin (host proteins normalized together over all times, phage proteins normalized by time-point). Limma was used to compare samples to the uninfected time-points, to compare genotypes at matched time-points, and to compare time-points across a specific phage infection. The model design included the phage genotype and time-point status for phosphoproteomics, or considering sample type (phage genotype + time-point) and replicate status for proteomics. The linear model was fitted using limma’s lmFit() function and a moderated t-statistic was computed using limma’s eBayes() function. P-values were adjusted for multiple testing using the Benjamini-Hochberg (BH) procedure implemented in limma. Proteins were considered to be statistically significantly changing if they had adjusted p-values <0.05 and absolute log2FC >log2(1.5). Phosphopeptides were considered to be statistically changing if they had adjusted p-values <0.05 and an absolute log2FC >1.

For the Retron-Eco9 and DarTG1 construct expression experiments, vsn-normalization was performed without grouping of samples/proteins. For the T7K shut-off domain solubility analysis, no vsn-normalization was performed. In both experiments, statistical significance was assessed using unpaired t-tests, using the stat_compare_means() function of the ggpubr R-package.

### Data processing and statistical analysis for stoichiometry experiment

For the stoichiometry analysis, only peptides that spanned residues previously found to be phosphorylated in the time-course phosphoproteomics experiment were considered. The following PSMs were not considered for downstream analysis: (i) PSMs without complete TMT labelling (i.e. without peptide N-terminal labelling), (ii) PSMs modified with S/T/Y phosphorylation or M oxidation variable modifications, (iii) PSMs mapping to proteins identified as differentially abundant in the time-course protein expression analysis (adjusted p-value <0.05, fold-change <0.5 or >1.5 at T=10 min, wild-type T7 vs Δ*0.7*).

Two limma analyses were performed, and both analyzed summed peptide intensities that were vsn-normalized without any sample/protein grouping. The first was performed to calculate and assess the quality of phosphatase/control ratios for each phage genotype (wild-type T7, Δ*0.7*) and each of three biological replicates, by inputting summed vsn-normalized TMT reporter intensities and comparing phosphatase vs control samples. It considered the sample (WT-phosphatase, WT-control, Δ*0.7*-phosphatase, Δ*0.7*-control) and replicate status in the model design. Differential analysis with limma was performed as described above. We considered some peptides to have poor phosphatase/control ratios when they were found in this analysis to be either significantly down-regulated (limma, adjusted p-value < 0.05, logFC < 0) or the log2FC was low (logFC < −0.2, no significance requirement). These were excluded from the subsequent analysis. The estimated stoichiometry value for each genotype was calculated from these ratios using the following calculation: 100*(1 - 1/2^log2FC). The T7K-mediated stoichiometry was calculated by subtracting the estimated stoichiometry of the Δ*0.7* from the wild-type T7 sample.

The second limma analysis was performed to determine differential stoichiometry i.e. stoichiometry deposited by the T7K. It was assessed by inputting manually-calculated phosphatase/control ratios for each genotype and replicate for the peptides that had quality phosphatase/control ratios (see above) in both samples (T=10 wild-type and T=10 Δ*0.7*). These were manually-calculated as: 2^summed vsn-normalized Phosphatase intensity / 2^summed vsn-normalized Control intensity. The limma model was constructed to consider the phage genotype (WT T7 or Δ*0.7*) and replicate status. Peptides were considered to be of “high” differential stoichiometry if they had adjusted p-values < 0.05 and log2FC > 1.5.

### Gene ontology analysis

To categorise phosphoproteins, goslim_prokaryotic_ribbon annotations were downloaded from QuickGO^109^ and annotated onto proteins. Proteins were assigned one of four representative GOSlim terms using the following ordered scheme - (i) transcription, translation (GOSlim terms: *ribosome biogenesis, translation, regulation of DNA-templated transcription, DNA-templated transcription*), (ii) metabolic processes (GOSlim terms: *metabolic process, primary metabolic process, lipid metabolic process,generation of precursor metabolites and energy, amino acid metabolic process, DNA metabolic process, small molecule biosynthetic process*), (iii) other (GOSlim terms: *cell wall organization or biogenesis, detoxification, protein folding, transport, response to stimulus*), or (iv) unnannotated, if it could not be mapped to a GOSlim.

Gene ontology (GO) term analysis was performed using the STRINGdb R package (version 2.14.3) ^90^, initialized with the settings: species = 511145, version = 12.0, score threshold = 400. The get_enrichment() command (with defaults) was applied. For the non-phosphorylated proteome subset, host proteins (≥ 1 unique peptide identified in the parallel proteomics measurement) that were not found to have any uniquely-mapping T7K-dependent phosphopeptides were compared to all detected host proteins (≥ 1 uniquely mapped peptide identified in the parallel proteomics measurement) as background. For the stoichiometry analysis, host proteins mapping to high differential stoichiometry peptides were considered, comparing to all detected host proteins (≥ 1 uniquely mapped peptide identified in the same experiment) as background.

### Comparisons to published phosphorylation datasets

Phosphosites for *Escherichia coli* (K-12) were downloaded from the dbPSP2.0^44^ in June 2024. Mass spectra from several previously published phosphoproteomics studies of *E. coli* (PRIDE submissions PXD008369, PXD008921 and PXD008289) were reanalyzed as described above, with IDs harmonized by use of the same FASTA database.

The fraction of phosphorylated S, T and Y residues in different datasets was calculated per phosphorylated protein as the number of phosphorylated S, T and Y residues divided by the total number of S, T and Y residues present in the protein’s sequence. For the T7 dataset, either (i) all detected sites from the TMT time-course, or (ii) only sites sensitive to T7K were considered. The phosphorylation fraction was also calculated for the human proteome based on (i) all phosphosites reported in the PhosphoSitePlus database^88^, (ii) a comprehensive collection human phosphosites measured by MS curated in a recent stringent meta-analysis^85^ and (iii) a single experimental dataset (using the same phosphoenrichment protocol as performed in this study) comprising phosphosites which were sensitive to EGF stimulation in HEK293F cells^86^. All distributions were compared to the distribution of T7-sensitive set of proteins using unpaired two-samples Wilcoxon signed-rank tests, which are implemented in the stat_compare_means() function of the ggpubr R-package.

### Phylogeny analysis

Amino acid sequences belonging to the “serine-threonine kinase” phrog 2828 (N=51, including T7K protein) were downloaded from the PHROG database (annotation release v4)^49^. The sequence of a recently reported homologous protein kinase JSS1_004^21^ from the *Salmonella* phage JSS1 (GenBank: OP185508.1) was also downloaded and considered. Jalview (version 2.11.41.1)^110^ was used to align the proteins belonging to the kinase phrog_2828 and JSS1_004 using the MAFFT webservice^111^. A phylogenetic tree was then generated on the multiple sequence alignment using the inbuilt Jalview function (neighbour joining, BLOSUM62 settings), and the iTOL tool^112^ used to visualize and annotate the resulting tree.

### Sequence motif analysis

For the analysis of T7K kinase activity motifs, the 10 amino-acid residues surrounding each side of a phosphorylated residue (only from sites with PTMProphet localisation score ≥ 0.75) were extracted and used to generate sequence logo plots using the ggseqlogo R package (version 0.2) ^113^.

### Protein structure analysis

Phage T7 protein structures were predicted by subjecting the Swissprot phage T7 proteome (UP000000840) sequences to AlphaFold3^59^ modelling via the AlphaFold Server. The PyMOL Molecular Graphics System (version 3.03 Schrödinger, LLC) was used to calculate the relative per-residue solvent accessible surface area on the top ranked model of each protein using the get_sasa_relative command.

FoldSeek^48^ structural homology searches were performed through the FoldSeek webserver, using the top-ranked AlphaFold3^59^ models generated for the full-length T7K and for T7K kinase-domain only (residues 1-242), and searching all available databases in 3Di/AA mode.

For the analysis of the T7K AlphaFold3 protein structure, PyMOL was first used to convert from .cif to .pdb format. This .pdb file was submitted to the GraphBind server^60^, with all ligand-specific options (including DNA and RNA) considered. Residues reported to be DNA-or RNA-binding are those that passed GraphBind’s internal binary thresholds.

### Phage infection assays of *E. coli* K-12 +/- various plasmids

To test a wide range of phage MOIs, serial dilutions of phage lysates (10 μl final volume) were added in round-bottom 96-well plates. We based MOI calculations on the assumption that an optical density of 1 corresponds to 1·10^9^ bacterial cells. Plates containing the phages were pre-warmed at 37°C for 1 h before adding the cells. *E. coli* BW25113 strains carrying plasmids encoding defense systems (with native promoters or with the inducible pBAD promoter; **Supplementary Table 5**), or a control plasmid, were diluted to 1:2,000 from overnight cultures and grown at 37°C. Strains with a plasmid with a native promoter were grown until OD600 = 0.05 (1 cm pathlength); strains with a plasmid with an inducible pBAD promoter were grown until OD600 = 0.6 (1 cm pathlength) in arabinose-containing medium (0.2% L-arabinose) as the pBAD promoter is catabolite-repressed in LB during exponential growth. Note that for Retron-Eco9 (**Fig. 3a and Extended Data Fig. 7e**), infection at OD600 = 0.05 (1 cm pathlength) led to inconsistent results, possibly because the culture entered stationary phase before the phage managed to collapse the bacterial population. We therefore started to dilute the cells 1:10 when they reached OD600 = 0.05, infecting the cells at OD600 = 0.005, which resulted in consistent phenotypes and effect sizes. Then, 90 μl of culture was added to the 96-well plates that contained phages and the plates were sealed with breathable membranes (Breathe-Easy). We kept several LB-filled wells in each experiment for background subtraction and to check for potential contamination. Plates were then incubated with constant shaking at 37°C in a microplate reader, and OD600 was measured every 10 min to acquire growth curves (either in a Biotek Synergy HT or Biotek synergy H1). Note that the OD600 values presented throughout all figures are from a microplate reader and were not pathlength-corrected.

### Phage infection assays of natural isolate library

The natural isolate strain collection (536 unique strains, 558 strains in total; **Supplementary Table 7**) is organized on 6 different 96-well plates. We reorganized some strains so that each plate contained at least 2 empty wells to check for cross-contamination and to use for background subtraction. The whole collection was screened in 4 batches.

Phage lysate stocks were prepared to account for final MOIs of 5, 0.01, 0.001, 0.0001 and 0, determined for the reference strain BW25113, and assuming that an OD600 of 1 corresponds to 10^9^ bacterial cells. We diluted the overnight cultures 1:2,000 and grew the strains at 37°C and 800 rpm in deep 96-well plates until the first strains reached OD600 = 0.1 (1 cm pathlength). Note that the strains grow at different rates, which makes it impossible to infect them all at the exact same OD600 = 0.1 (1 cm pathlength). Therefore, the exact phage-bacteria ratio will vary for each strain, but we tried to account for that by applying a range of MOIs. We then pipetted 90 µl of the bacterial cultures into 9 different round-bottomed 96-well plates, and added 10 µl of phage lysate on top (a single MOI per plate). Plates were sealed with breathable membranes (Breathe-Easy) and incubated without lids in a humidity-saturated incubator (Cytomat 2, Thermo Scientific) with continuous shaking. Plates were measured every 45 min in a Filtermax F5 multimode plate reader (Molecular Devices).

Out of the 558 strains, the uninfected control (MOI = 0) did not grow for 6 strains (ECOR-70, MG1655, ATCC 8739, S17 plate1, DH1, DH1 (E381)), so they were removed from the analysis. Strains with sequences not matching the expected genotype were also removed from the analysis (all strains with a/b/c/x as a suffix in their name, **Supplementary Table 7)**; note that 17 strains are present in replicates (same strain names) within the collection and we added the plate number behind the name to distinguish these in the analysis (examples in **Extended Data Fig. 10**). One strain (NR-2654) appeared twice on the same plate, so we added _1 and _2 to the name, respectively. Three strains (IAI01, S17, and NR-2654) had replicates with different strains IDs (‘NT numbers’), and we counted them only once among the unique strains. All replicates fell in the same categories (uninfected, infected without evidence of defense, infected with evidence of defense), except for three (MP1, ECOR-09, ECOR-10). For these strains, we removed the replicate from the ‘uninfected’ category, as MP1 was validated from an independent glycerol stock to be infected **(Extended Data Fig. 11)**, and ECOR-09 and ECOR-10 were slowly growing in the replicate that showed infection **(Extended Data Fig. 10b)**, so the other replicate was probably contaminated. For the analysis we therefore assessed 513 unique strains (529 strains in total). Four other strains (BL21(DE3), DH5α, ECOR-09 plate5, ECOR-10 plate5) grew very slowly and the integration time for the AUC calculation (see below) was therefore increased from 6 to 10 h **(Extended Data Fig. 10b)**. The whole collection was screened in one replicate, and all 22 strains showing infection and evidence of defense **(Extended Data Fig. 10c)** were repeated from an independent glycerol stock and with higher MOI resolution (validations) in at least two biological replicates, i.e. starting from two independent colonies on a plate **(Extended Data Fig. 11)**.

### Data processing of phage infection assays

All the data from plate reader assays were processed and plotted using Matlab R2017b (The MathWorks). The data were background-subtracted with one background value (minimum measured value across each plate and time-points) and plotted over time. For the natural isolates screen we subtracted the background at each time-point with the minimum measured value across all plates of a batch at the time point. The area under the curve (AUC) was determined under the linear growth curve from 0 h until 6 h of measurement using the function trapz. AUCs were normalized to the AUC of the uninfected control (MOI = 0) of the same strain (normalized AUC). To estimate the effect size for a biological replicate of a strain, we interpolated the MOI at an AUC of 0.5 on the log10 scale for both T7 WT and Δ*0.7*, using the Matlab function interp1. Then we took the inverse logarithm base 10 of the difference (Δ*0.7* minus WT). As the effect size strongly depends on well-matched phage titers of T7 WT and Δ*0.7*, we only considered a mean effect size of > 5-fold or < 0.2-fold as a hit **(Fig. 4b and Extended Data Fig. 11c)**.

### Bacterial genome analyses

Genomic nucleotide sequences for the majority of the *E. coli* natural isolates were downloaded from the ECOREF^72^ resource’s website (https://evocellnet.github.io/ecoref/). The genomes of several strains (IAI09, IAI41, Perfeito-Clone-6, H10407, HM-341) were additionally sequenced using long-read sequencing. Overnight cultures of strains grown at 37°C, shaking, were used for DNA extraction with the DNA mini-prep kit (ZymoBIOMICS). For library preparation, 1.2 μg of DNA for each sample was taken into fragmentation using Megaruptor 3.0 (Diagenode) with a speed of 31 and a final volume of 100 μl. Then 1X clean-up with SMRTBell beads (PacBio) was performed. All recovered material was used as an input for library preparation using SMRTBell 3.0 chemistry (PacBio) and following the manufacturer’s instructions. Final libraries were quantified by Qubit and Femto Pulse system was used to assess the fragment distribution. Libraries were then pooled equimolarly and loaded in the Sequel IIe instrument at 100 pM with 30 h movie time. The resulting PacBio reads were assembled into high-quality, complete genomes using Flye^114^ (version 2.9-b1768) using standard parameters.

To annotate known bacterial anti-phage defense systems in natural isolate strain genomes, nucleotide sequences were either (1) inputted directly into the PADLOC python program (version 2.0.0, using defaults)^74^ using PADLOC-DB version 2.0.0, or (2) first translated into a protein FASTA file using the Prodigal python package (version 2.6.3, using defaults)^115^, and then inputted into the DefenseFinder python package (version 1.2.2, using defaults)^1^ using DefenseFinder models (version 1.2.3)^73^ and CasFinder models (version 3.1.0)^116^.

---

**Extended Data Figure 1:**
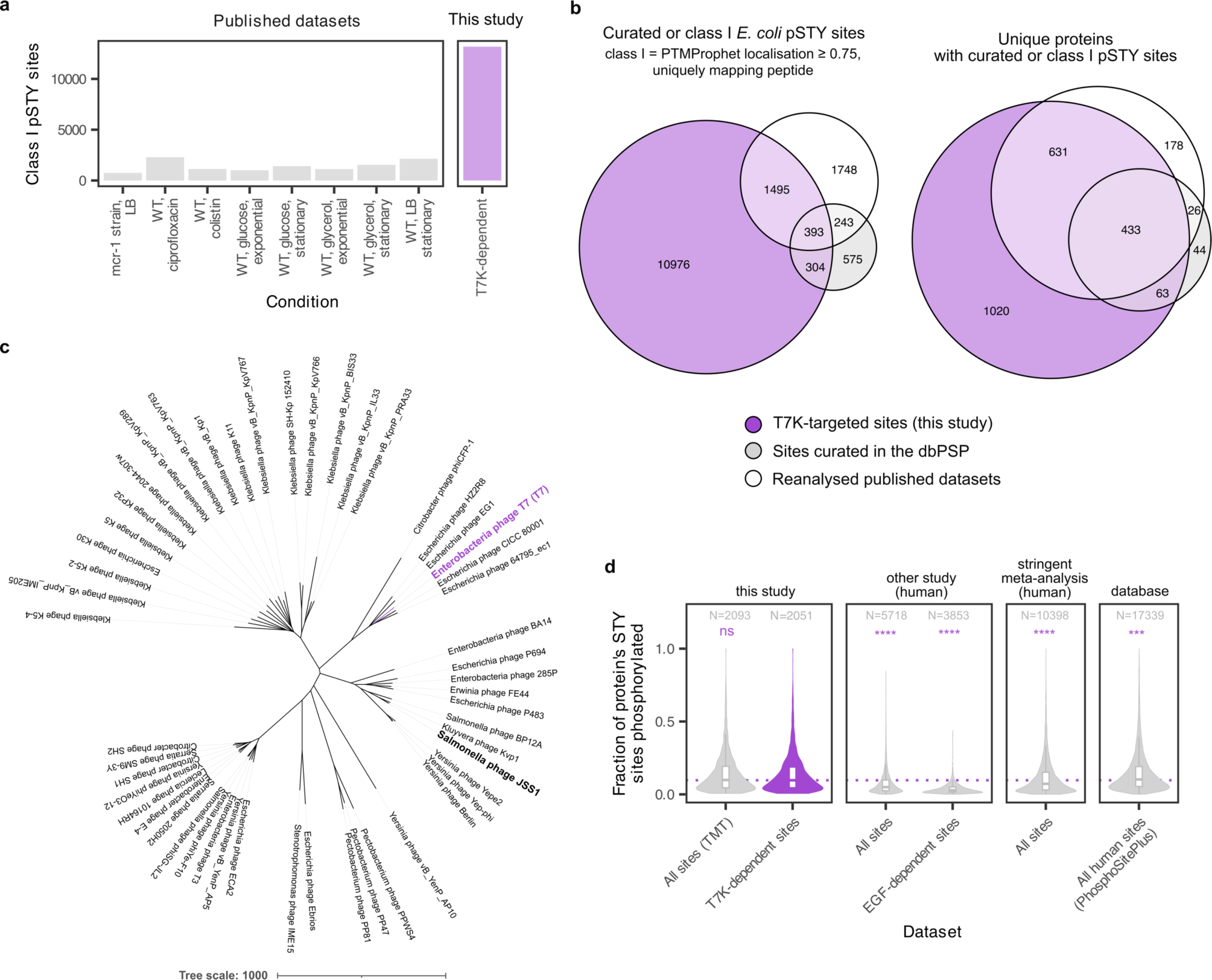
T7K is a hyper-promiscuous protein kinase. **(a)** The number of well-localized phosphosites on serine (S), threonine (T) and tyrosine (Y) residues that were T7K-dependent compared to several other published phosphoproteomic datasets of various cellular perturbations of *E. coli*. These included *E. coli* (i) grown in either glycerol or glucose at exponential or stationary phase^42^, (ii) grown in minimal media^45^, or (iii) under antibiotic stress, with and without genetic resistance to the colistin antibiotic (mcr-1 mutant strain)^46^. All datasets were re-analyzed using the same MSFragger settings and FASTA database to facilitate comparison. T7K-dependent refers to sites from phosphopeptides significantly upregulated in wild-type T7 infection relative to uninfected, and also relative to the Δ*0.7* infection at a matched time-point (limma, FC > 2, adjusted p-value < 0.05). Class I sites have a high confidence of the modification’s localization within the peptide (PTMProphet^87^ localization score ≥ 0.75). **(b)** Overlap of phosphosites and phosphoproteins with relevant databases and the reanalyzed, previously published datasets. Class I phosphosites considered to be T7K-targeted were those that were host sites and were either (i) found uniquely in the wild-type T7 infection samples in label-free analysis described in Fig. 1b, or (ii) found to be T7K-dependent in the TMT time-course (see above). These sites and resulting phosphoproteins were compared to the curated *E. coli* sites in the bacterial phosphorylation database dbPSP^44^, and also to the bacterial mass-spectrometry based studies of the *E. coli* phosphoproteome under various perturbations which we reanalyzed as described above. **(c)** Phylogenetic tree of protein sequences relating to T7K (see Methods), labelled with their phage species of origin. These include phage kinase proteins in the same orthology group as curated by the PHROG database (phrog 2828, N=51 sequences) which includes T7K itself (bolded and purple). The JSS1_004 kinase protein (bolded) reported in ^21^ was also considered. **(d)** Extension of Fig. 1i, showing the per-phosphoprotein (S/T/Y) phosphorylation fraction across several sources of information on the human phosphoproteome. T7K-dependent sites (purple) are the same as described above. All sites (TMT) refers to all phosphosites identified and quantified in the TMT experiment (regardless of T7K-dependency). “other study (human)” refers to a recent phosphoproteomics study from our group investigating phosphoregulation in human HEK293F cells after EGF stimulation^86^, which used the same phosphoenrichment strategy. “All sites” refers to all identified and quantified sites in this study, whereas “EGF-sensitive sites” are those reported by the paper to be significantly induced by EGF stimulation, which is known to rapidly induce a wave of downstream phosphosignalling in the human cell. “stringent meta-analysis (human)” refers to all phosphosites reported in a recent meta-analysis and curation of the human phosphoproteome^85^, and “database” refers to all human sites listed on the PhosphoSitePlus database^88^. The distributions of the different sets were compared to the T7K-dependent set using unpaired two-sample Wilcoxon signed-rank tests (**** meaning p-value ≤ 0.0001, *** meaning p-value ≤ 0.001). Numbers indicate the number of phosphoproteins considered in each distribution. The boxplot shows the data distribution, with the box spanning the interquartile range (25th and 75th percentile) and the median indicated at the line. Whiskers extend to the smallest and largest values within 1.5 × IQR, while outliers are not plotted as they are encompassed by the corresponding volcano.

**Extended Data Figure 2:**
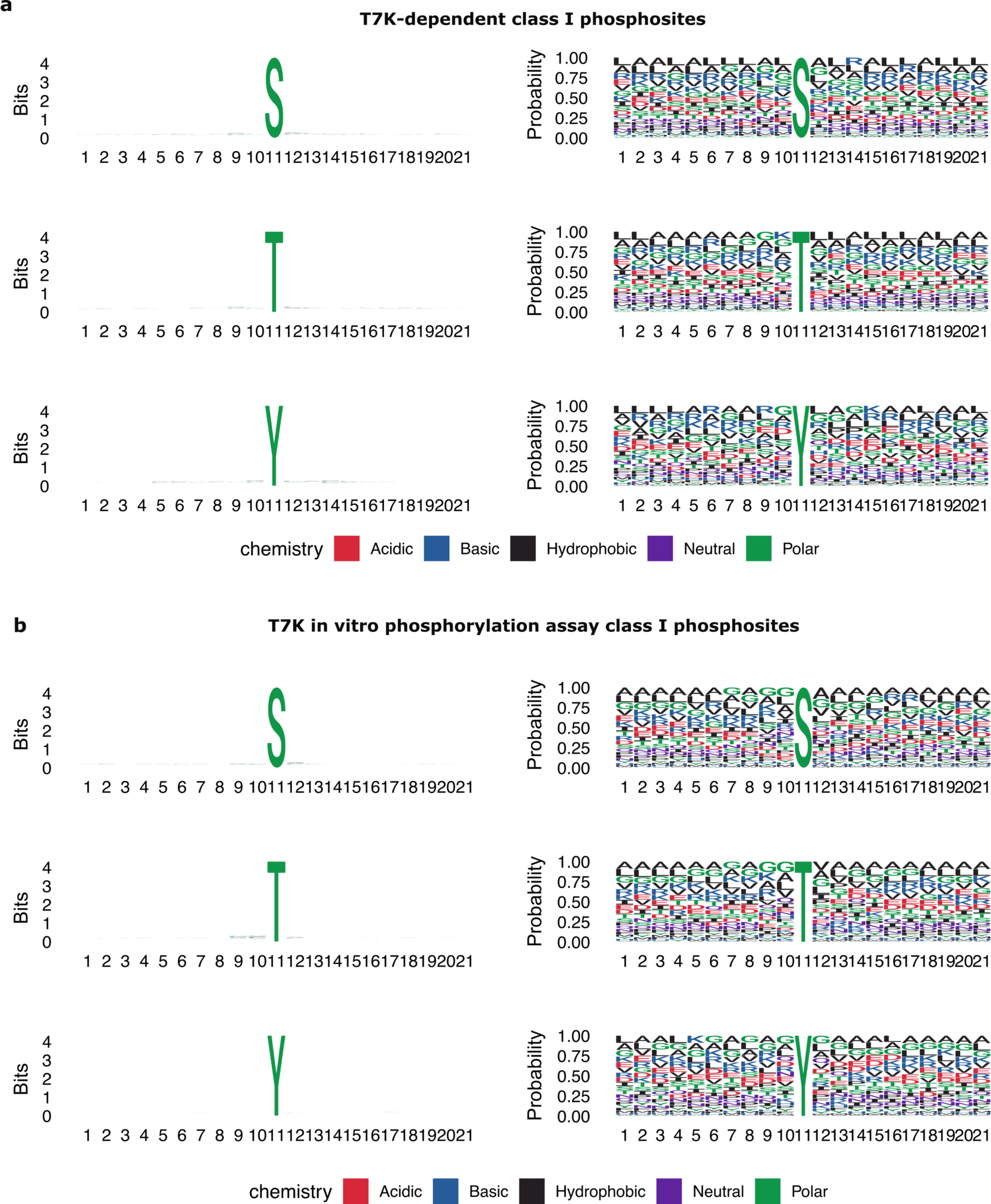
Phosphosite motif analysis. Motif analysis of phosphosites from class I phosphopeptides (PTMProphet localisation ≥ 0.75, uniquely mapped to a single protein) from either **(a)** T7K-dependent phosphopeptides identified in the TMT time-course (Fig. 1e) or **(b)** from phosphopeptides identified in the in vitro phosphorylation assay using the purified T7K kinase domain (Fig. 1c).

**Extended Data Figure 3:**
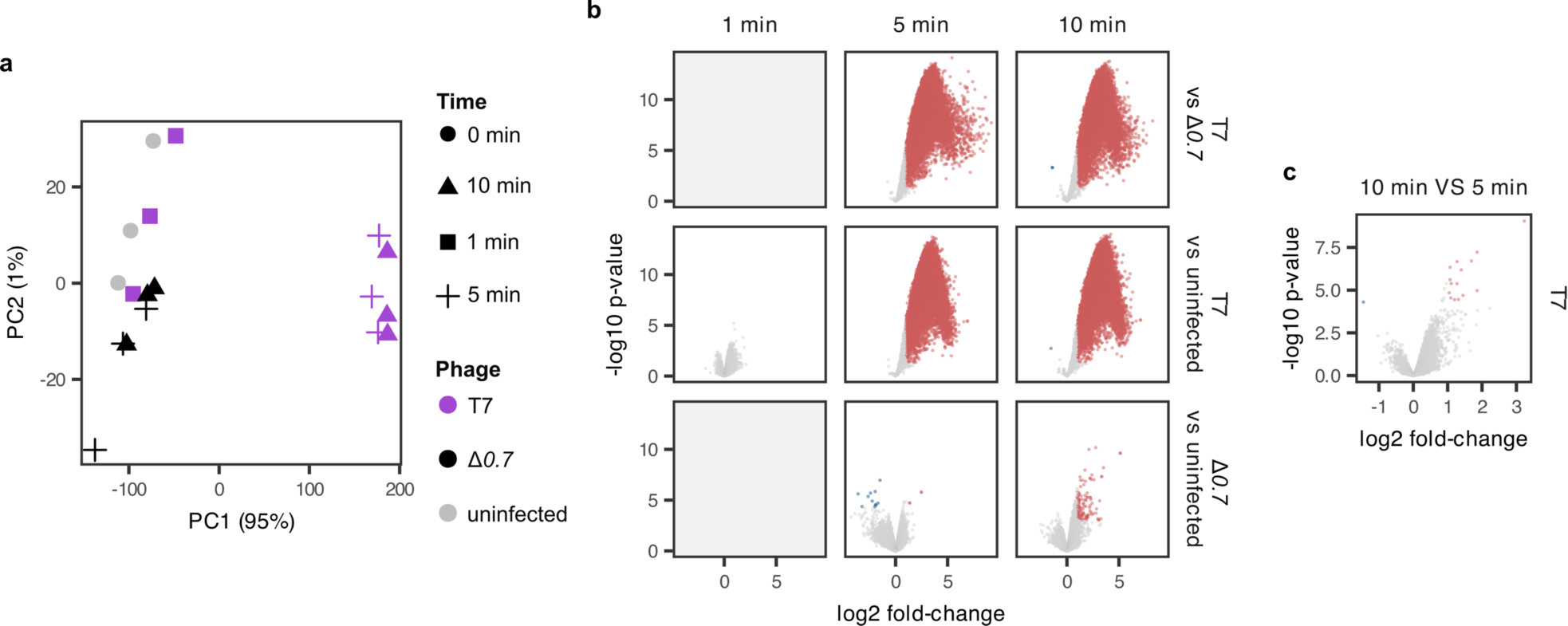
Statistical analyses relating to the analysis of T7 infection time-course phosphoproteomics dataset. **(a)** Principal component analysis (showing top-2 principal components) of samples based on vsn-normalized^89^ phosphopeptides with full quantification in the TMT-multiplexed time-course analysis of wild-type T7 and Δ*0.7* infections, across three biological replicates (Fig. 1e). **(b)** Volcano plots showing the results of limma differential phosphopeptide abundance analysis for several comparisons of wild-type or Δ*0.7* infections to the uninfected sample, or to each other at matched time-points. Red and blue dots show phosphopeptides with significance (limma, adjusted p-value < 0.05, 2-fold change) with upregulated sites as red and downregulated sites as blue. Grey facets show comparisons that were not possible (comparison against 1 minute infections involving Δ*0.7*) due to the TMT-multiplexing design constraints. **(c)** Volcano plot showing an additional comparison between 5 and 10 minutes of wild-type T7 infection, illustrating minimal changes to the phosphorylation levels which likely reflects deactivation of the kinase activity by autophosphorylation.

**Extended Data Figure 4:**
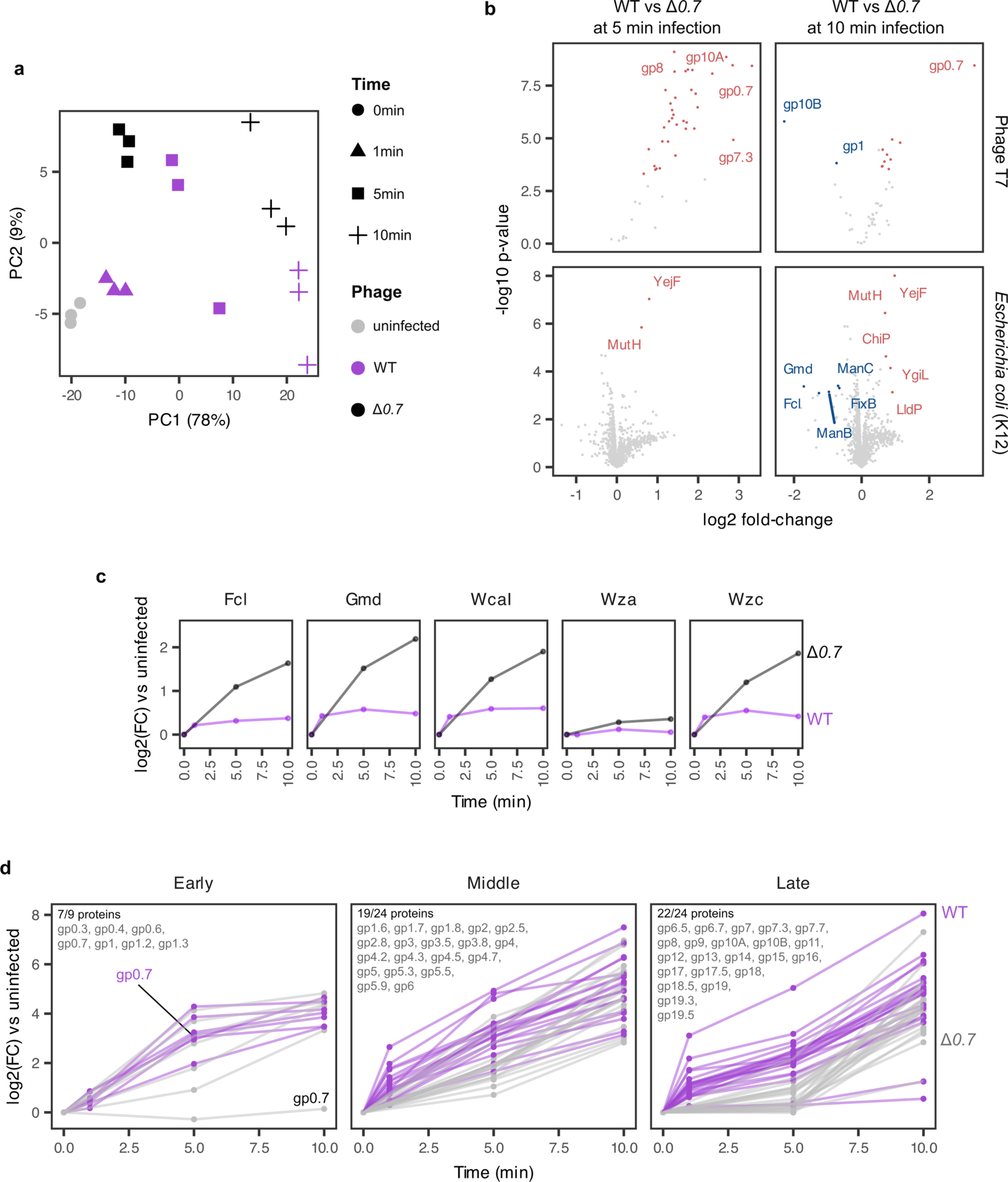
Analyses relating to the analysis of T7 infection time-course proteomics dataset. **(a)** Principal component analysis (showing top-2 principal components) of samples based on vsn-normalized^89^ protein intensities with full quantification in the TMT-multiplexed time-course analysis of wild-type T7 and Δ*0.7* infections, across three biological replicates (Fig. 1e). **(b)** Volcano plots showing the results of limma differential protein abundance analysis for wild-type T7 infection compared to Δ*0.7* infection at 5 minutes and 10 minutes. Red and blue dots show proteins with significance (limma, adjusted p-value < 0.05, 1.5-fold change) with upregulated sites as red and downregulated sites as blue, and proteins are grouped by their organismal source. N = 48 phage proteins for the top facet, and N = 2,919 host proteins in the bottom facet for each volcano plot. **(c)** Selected time-course measurements (log2FC of infected sample at given time-point vs uninfected) for host proteins regulated within the *wza* operon. Purple profiles show the protein’s expression pattern upon infection with wild-type T7 and black in Δ*0.7* infection. **(d)** Time-course measurements for phage T7 proteins classified by T7 gene class. The profiles for T7K (gp0.7) are indicated. The top left corner of each sub-graph shows the numbers and (gene product) names of phage proteins within each category that were successfully quantitatively monitored.

**Extended Data Figure 5:**
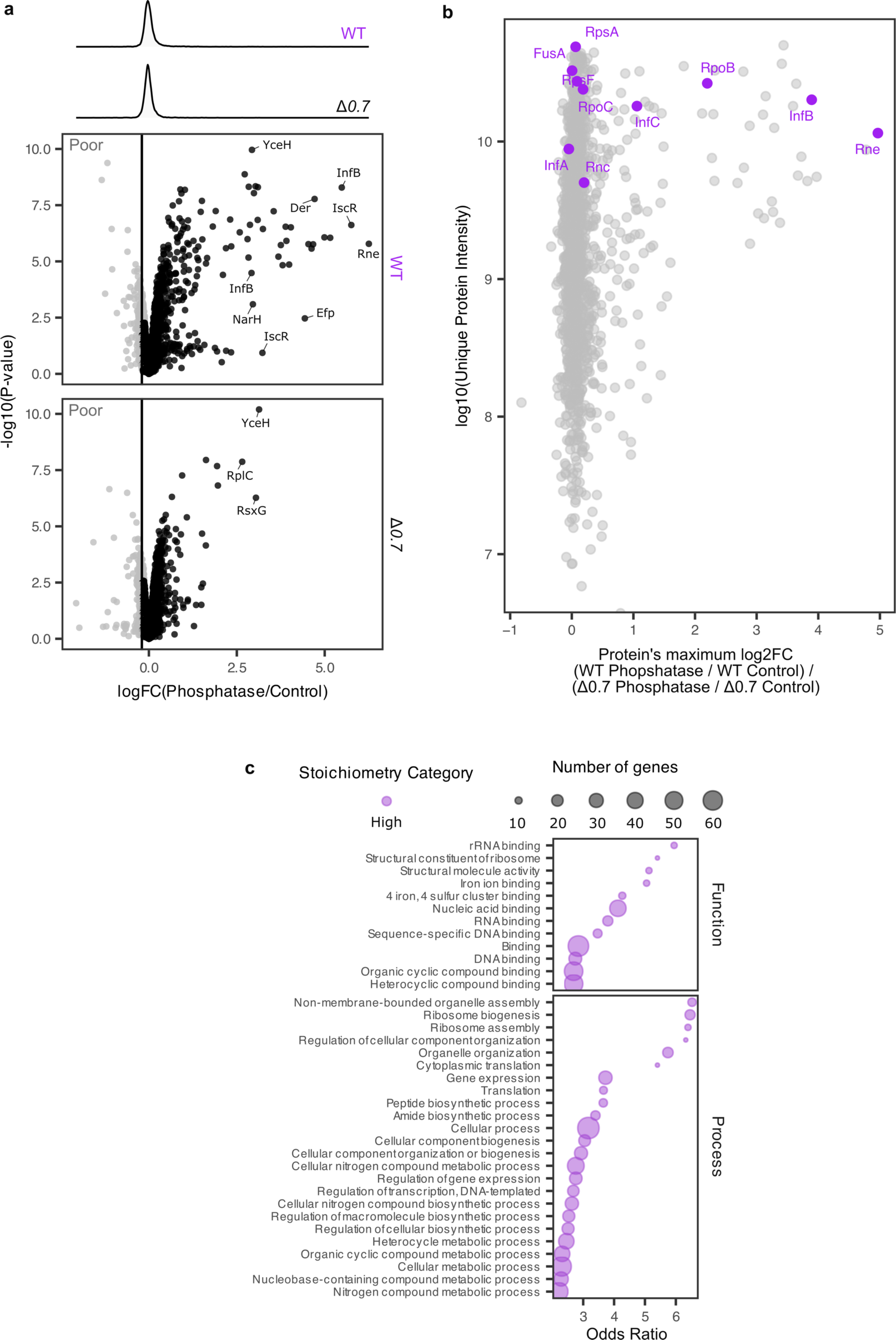
Estimating phosphorylation stoichiometries. **(a)** The Phosphatase/Control ratios of peptides considered for the estimation of phosphorylation stoichiometries (N = 4,371 peptides). These were (non-phosphorylated) peptides with residues previously identified to be phosphorylated in the TMT time-course phosphoproteomics dataset (Fig. 1e). Stoichiometries were estimated using three biological replicate samples of the T=10 minutes infection time-points, and as such only phosphosites on peptides representing proteins that were not found to significantly differ between WT T7 and Δ*0.7* infection at 10 minutes in the protein expression analysis (Extended Data Fig. 4b) were considered. We excluded “Poor” Phosphatase/Control ratios (grey dots) from downstream differential stoichiometry analyses using limma (Fig. 2b) when the Phosphatase/Control ratio was either significantly down-regulated (limma, adjusted p-value < 0.05, log2 fold-change (FC) < 0) or the log2FC was low (logFC < −0.2, no significance requirement). The log2FC = −0.2 position is indicated. This represented the minority of ratios (see corresponding density histogram for each sample, top of the graph), but removed peptides that had clear evidence of confounding technical factors (as the addition of phosphatase should not result in a decrease in signal, in theory). Both ratios were categorised as poor if they were found to be poor in at least one of the conditions (wild-type T7 or Δ*0.7*). Several peptides with high stoichiometries are labelled with the name of the protein they belong to. **(b)** The maximum differential stoichiometry log2FC values for each protein compared to its abundance (unique intensity) (N = 1,538 proteins), where maximum refers to the highest log2FC obtained for any phosphopeptide mapping to the protein with quality ratios calculated from (a). Previously reported host substrates are indicated in purple and labelled with their protein names. **(c)** Full results of GO-term analysis on high stoichiometry phosphoproteins, which includes the data presented in Fig. 2d. Enrichment analysis was performed on *E. coli* proteins with peptides showing evidence of having **“**high” differential stoichiometry (see Fig. 1b). All GO-terms have a false discovery rate ≤ 0.05 (Benjamini-Hochberg corrected, using the STRING-db R package^90^), and the expressed *E. coli* proteome was used as background (all proteins identified by unique peptides in the proteomic experiment monitoring wild-type T7 and Δ*0.7* infections).

**Extended Data Figure 6:**
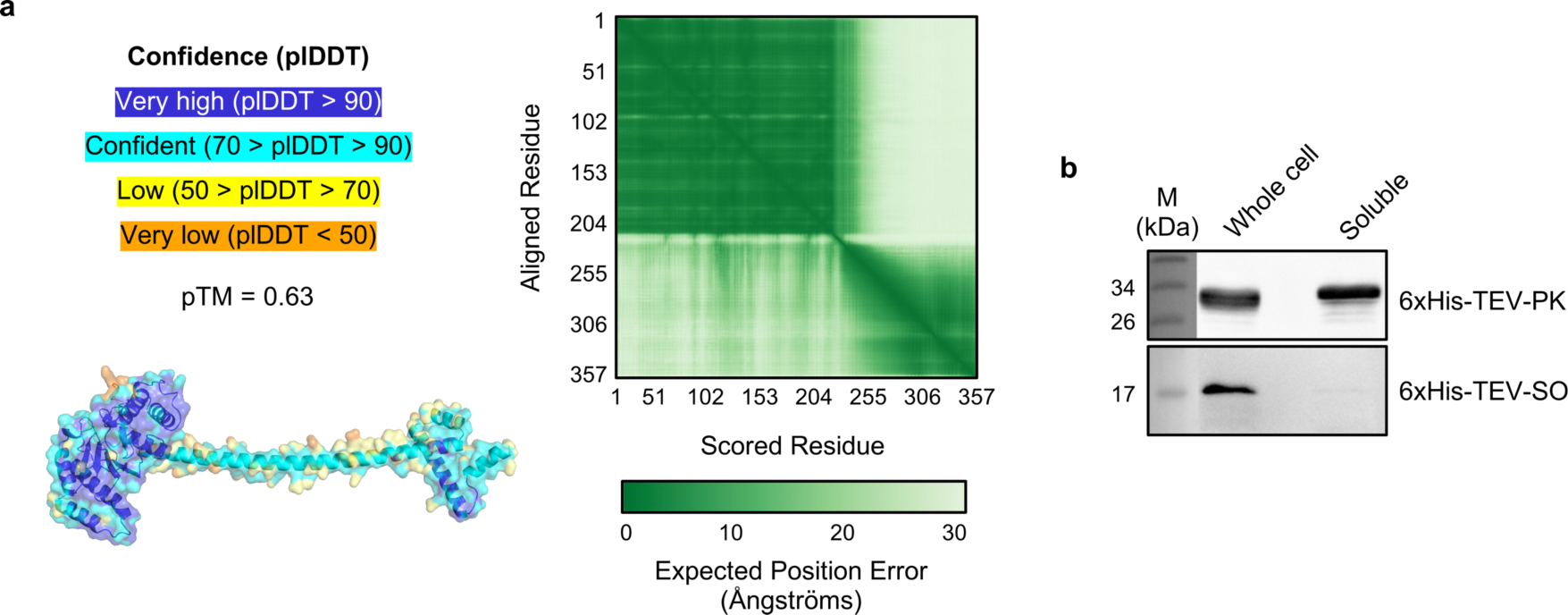
AlphaFold structural prediction and purification of T7K domains. **(a)** Prediction confidence scores for the top-ranked AlphaFold3^59^ structural prediction of T7K. Residues are coloured by their residue-level (predicted local distance difference test) plDDT score confidence category, with the structure illustrated from N-to C-terminus, matching the depictions of T7K in Fig. 2e. The residue-residue alignment confidence (PAE) matrix and predicted template modelling (pTM) score are also shown. **(b)** Western Blot analysis of 6xHis-TEV-PK and 6xHis-TEV-SO before and after cell fractionation. Proteins were expressed in *E. coli* BL21-AI cells 6xHis-TEV-PK and 6xHis-TEV-SO. Whole cell lysates or soluble cell extracts were loaded on 15% polyacrylamide and proteins resolved by SDS-PAGE, then transferred to PVDF. Membranes were probed with anti-His antibodies (Invitrogen) to detect proteins of interest.

**Extended Data Figure 7:**
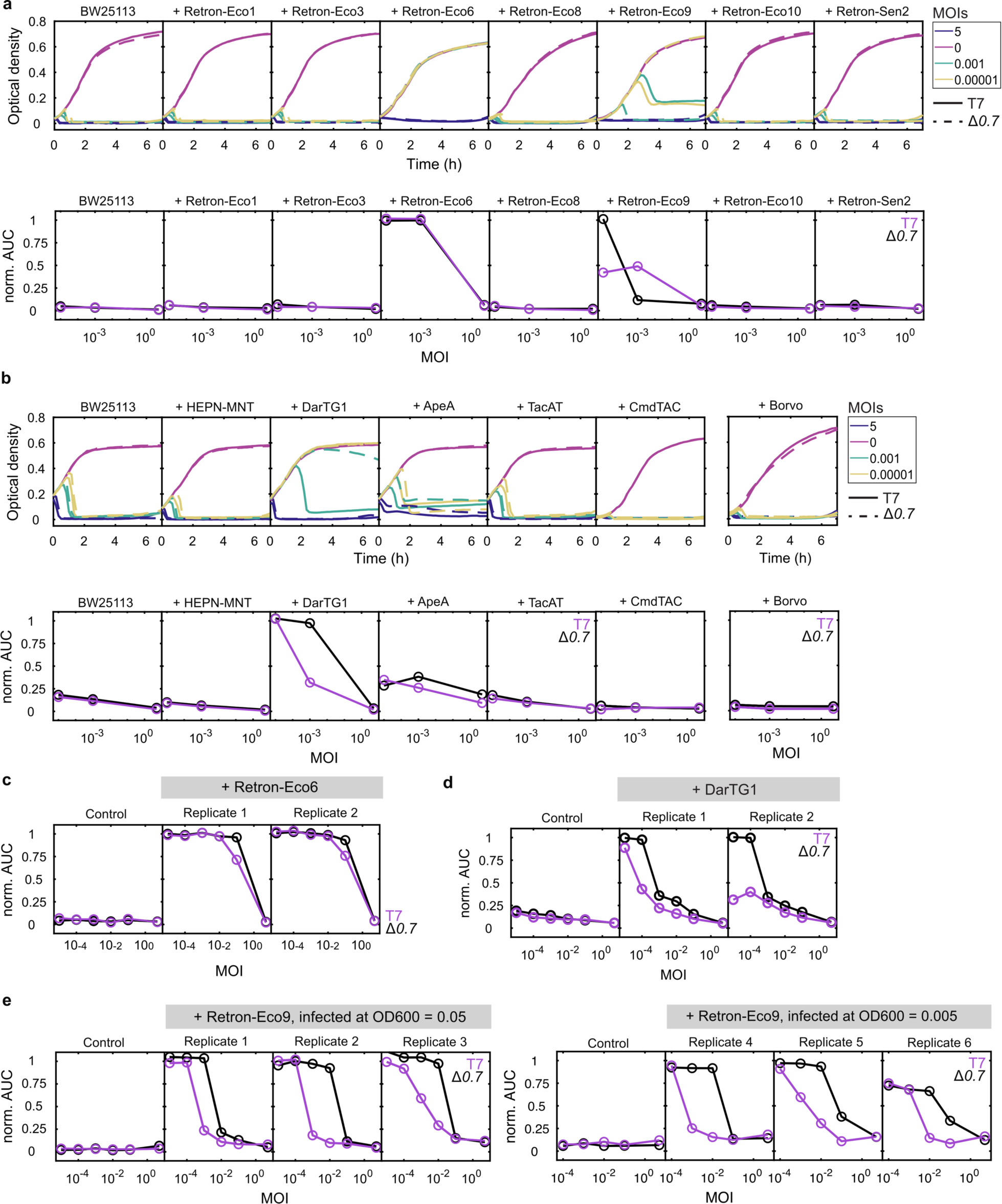
T7K weakens the activity of Retron-Eco9 and DarTG1. **(a)** Defense against T7 WT and Δ*0.7* by msDNA-containing retrons under native promoters. Upper panel: bacterial growth of *E. coli* BW25113 +/- the indicated defense systems over the course of T7 infection measured by optical density (600 nm) in a microplate reader. Lower panel: Normalized area under the curve (norm. AUC), normalized to the uninfected sample for all growth curves of the control BW25113 +/- defense systems infected with T7 (purple lines) or Δ*0.7* (black lines) at the indicated MOIs (empty circles). Data is the same as in the upper panel. **(b)** Defense against T7 by DNA/RNA-binding systems (left-side plots) and one CHAT-protease domain-containing system (Borvo; right-side plot). Upper and lower panels as in (a). **(c-e)** Norm. AUC graphs for all growth curves of the control BW25113 with empty plasmid or a Retron-Eco6 plasmid (**c**), a DarTG1 plasmid (**d**) and a Retron-Eco9 plasmid (**e**). All biological replicates shown - for DarTG1 and Eco9 biological replicates 1 and 2, respectively, are the same as in Fig. 3b. For Eco9, infecting at an earlier growth stage (OD600 = 0.005 vs. 0.05, measured with a pathlength = 1cm) made the effect size of T7K-sensitivity more consistent.

**Extended Data Figure 8:**
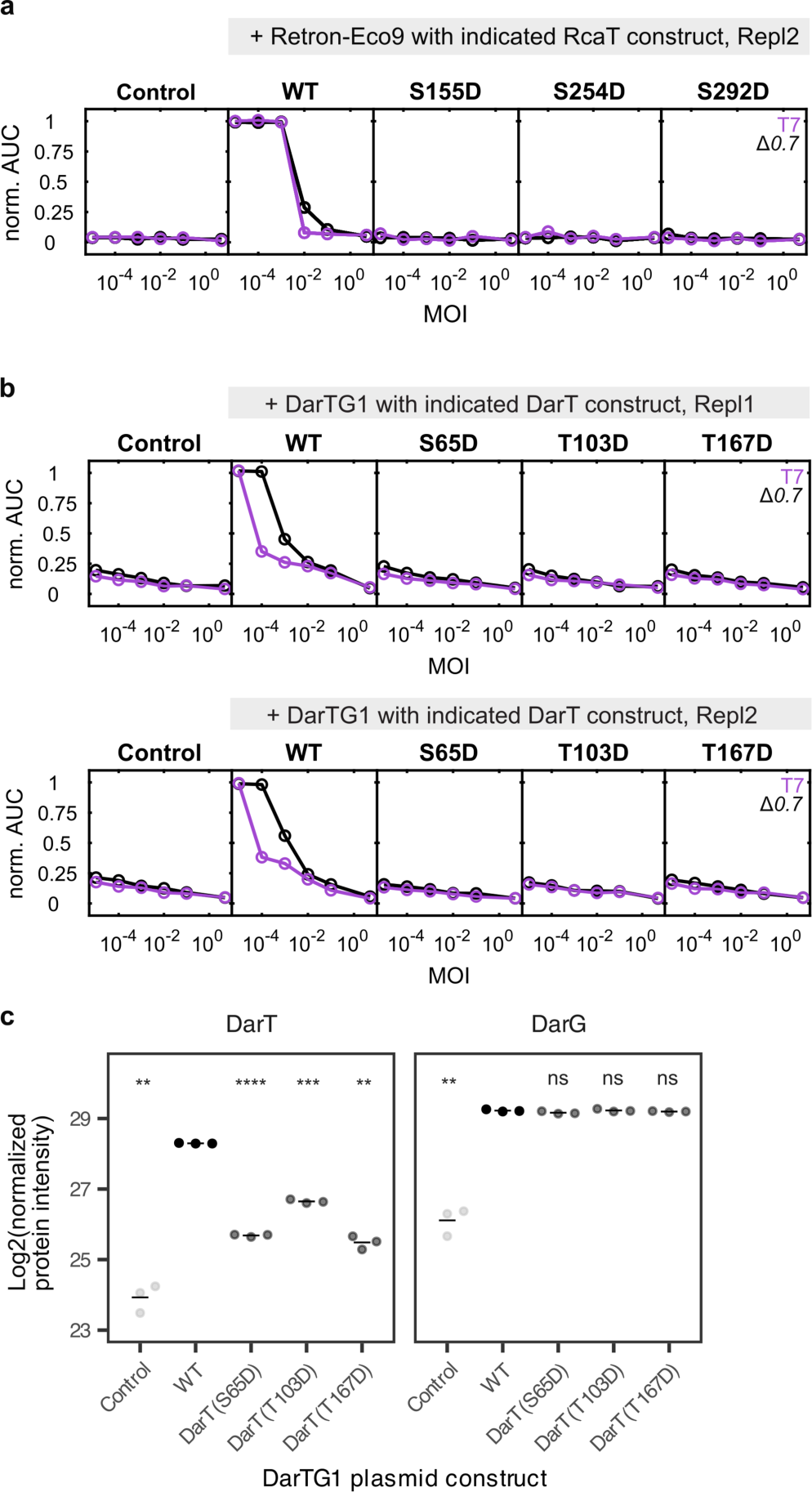
Phosphomimetic mutants of Retron-Eco9 and DarTG1. **(a)** Biological replicate of Fig. 3e. Normalized area under the curve (norm. AUC) for all growth curves of BW25113 with empty plasmid, Retron-Eco9 WT, or Retron-Eco9 RcaT mutants S155D, S245D, or S292D, infected with T7 (purple lines) or Δ*0.7* (black lines) at the indicated MOIs (circles), normalized to the uninfected sample. **(b)** Norm. AUC for all growth curves of BW25113 with empty plasmid or DarTG1 WT, or DarTG1 DarT mutants S65D, T103D, or T167D, infected with T7 (purple lines) or Δ*0.7* (black lines) at the indicated MOIs (circles), normalized to the uninfected sample. **(c)** Expression of the DarTG1 proteins DarT and DarG was investigated in *E. coli* K-12 transformed with either an empty plasmid (Control), DarTG1 in its wild-type form (WT), or with three phosphomimetic mutations of the DarT protein, in the absence of phage infection. The results of unpaired t-tests comparing expression levels in each construct to those in the WT DarTG1 construct are shown, with ns meaning non-significant (p-value > 0.05), * meaning p-value ≤ 0.05, ** meaning p-value < 0.01 and *** meaning p-value ≤ 0.001.

**Extended Data Figure 9:**
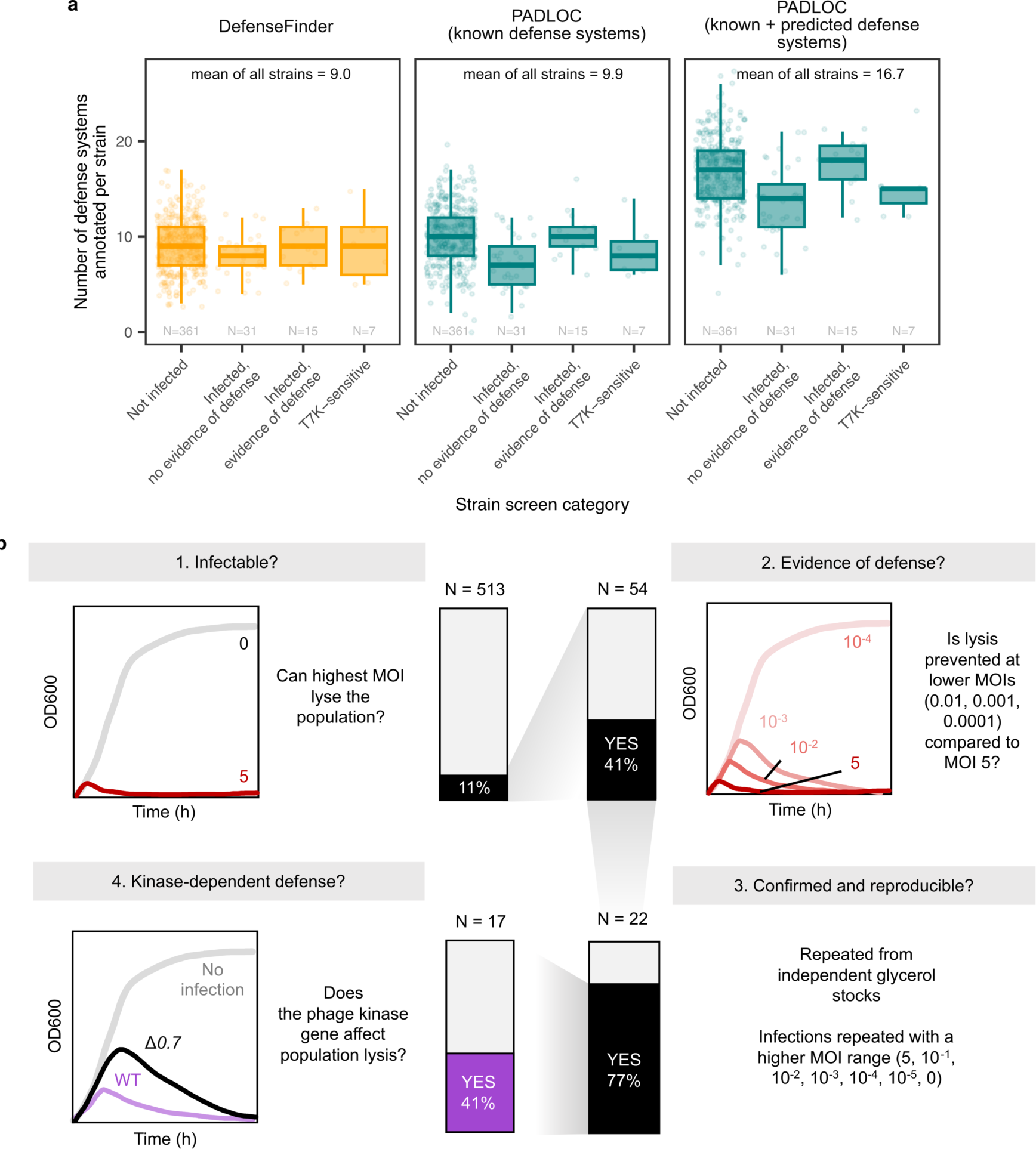
Overview of natural isolate library screen. **(a)** The number of defense systems detected within *E. coli* strains in the natural isolate collection which have sequenced genomes (N = 414), stratified according to the categories described in the following panel. Note: strains are categorized by their phenotypes in the initial screen, except for the fourth category which is made up of the 7 strains with *validated* T7K-sensitive defense, reported in Fig. 4b. Strains were annotated by the PADLOC^74^ and DefenseFinder^1^ tools. PADLOC also annotates phage defense candidate systems, with the prefix “PDC-”, which are not yet all experimentally validated to confer defense, so two distributions of the number of defense systems per strain for this tool are shown. The mean of the entire group is shown above the plot, and the mean and N (number of strains) for the subgroups are indicated on the boxplot. Boxplots are shown as in Extended Data Fig. 1e. **(b)** Schematic showing the categorization of strains in the natural isolate screen during the screen for strains that are sensitive to the presence of the T7K during infection by T7. N indicates the number of strains passing each filter, for example 54 strains were infectable by T7, and 22 T7-infectable strains had evidence of some existing defense against T7 infection. Results from these 22 strains (including those suggesting kinase-sensitive and kinase-insensitive defense) were then validated from independent single glycerol stocks and using a higher range of MOIs. Strains that passed this validation are the strains shown in Fig. 4b-c.

**Extended Data Figure 10:**
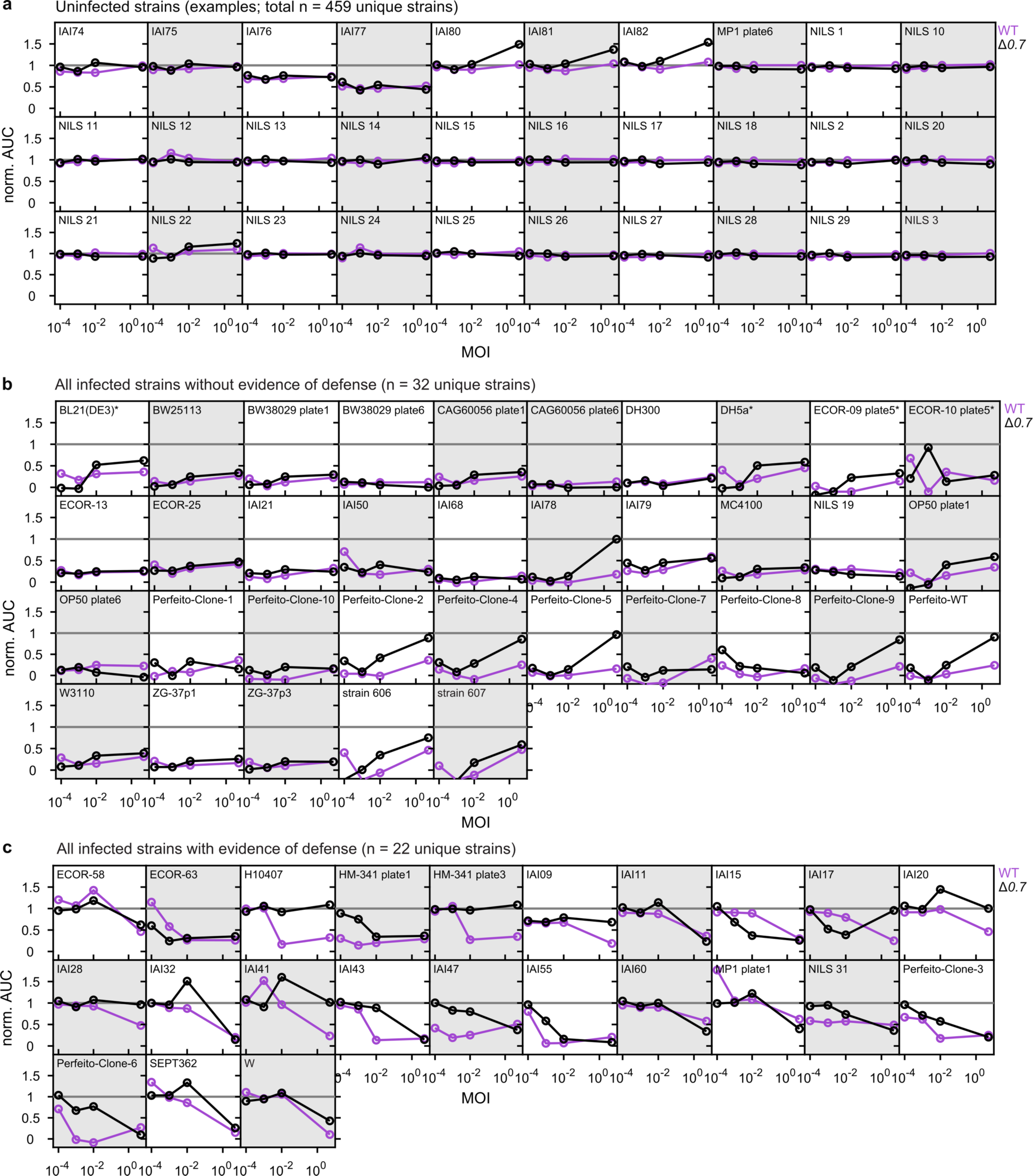
Natural isolate strains grouped into three categories: uninfected, infected without evidence of defense, and infected with evidence of defense. **(a)** Normalized area under the curve (norm. AUC) over multiplicity of infection (MOI) plot for 30 example strains that were not infected by T7 WT or Δ*0.7*. Their norm. AUC is constant over MOIs with a value of close to 1, meaning they grow as well as the uninfected control at all MOIs. Values may be consistently but slightly below 1 (e.g. strain IAI77) due to a technical reading issue in the uninfected control. **(b)** Norm. AUC over MOI plot for all strains that got infected by T7 WT or Δ*0.7*, but did not show any evidence of defense. Their norm. AUC is constant over all MOIs with a value close to 0, meaning that they do not grow at all, except in the uninfected control. For strains labeled with *, the integration time for norm. AUC was increased from 6h to 10 h, as they were growing very slowly. Values clearly above 0 are either due to technical problems or the development of resistance at later time-points. Strains present in replicates in the library have the same background color which alternates between white and grey. **(c)** Norm. AUC over MOI plot for all strains that were infected by T7 WT or Δ*0.7*, and had evidence of defense. Their norm. AUC decreases with increasing MOI, in either T7 WT or Δ*0.7*, meaning that they grew like the uninfected control at lower MOIs, but collapsed at the higher MOIs. All strains were infected at 5 MOIs (5, 10^-2^, 10^-3^, 10^-4^, and 0) with T7 WT and Δ*0.7*. Strains present in replicates in the library have the same background color which alternates between white and grey.

**Extended Data Figure 11.**
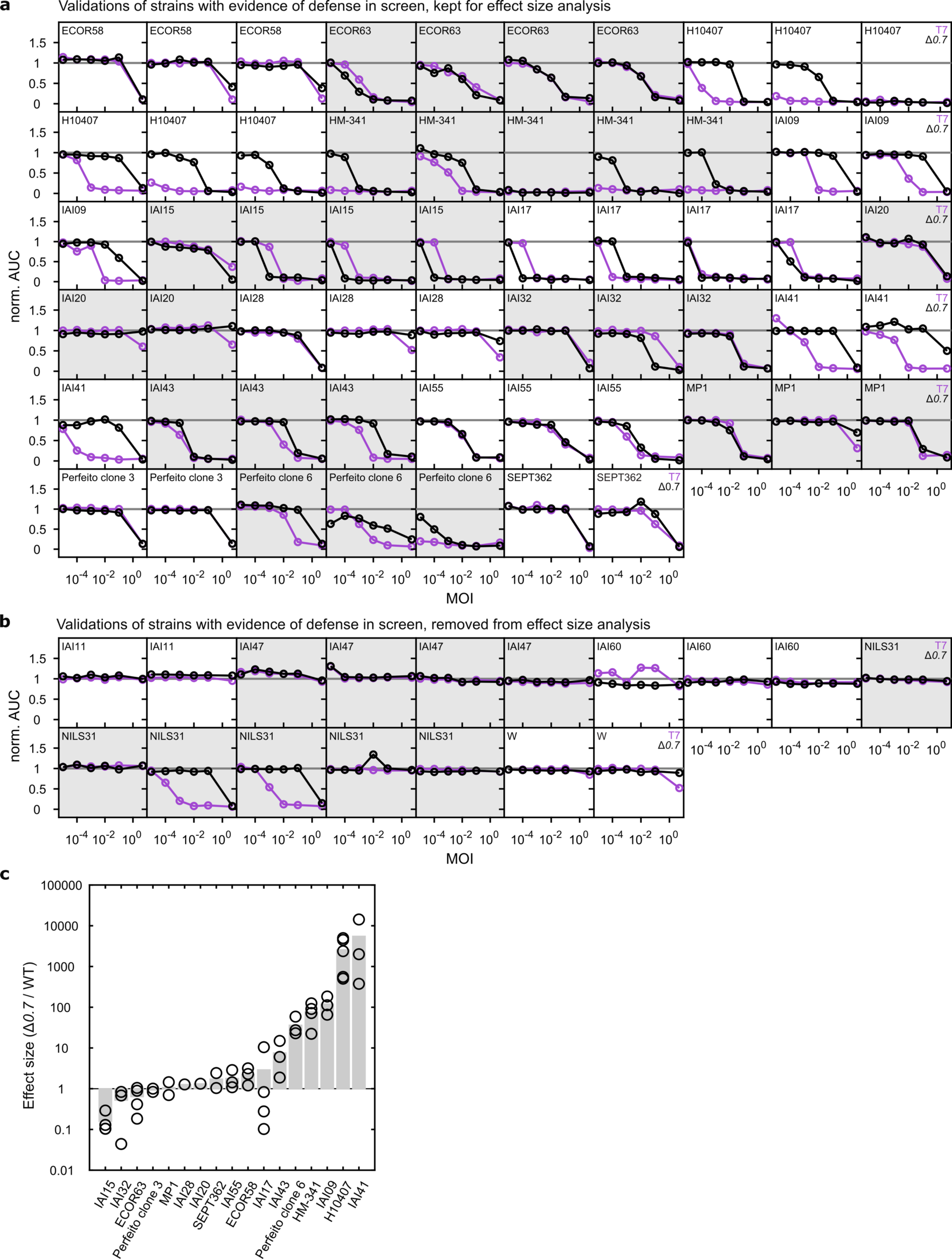
Validations of natural isolate strains that were infected by T7, but showed evidence of defense in the screen. **(a)** Normalized area under the curve (norm. AUC) over multiplicity of infection (MOI) plots for strains that were infected by T7, but showed evidence of defense (Extended Data Fig. 10c). All strains originate from individual glycerol stocks and were infected at seven MOIs (5, 10^-1^, 10^-2^, 10^-3^, 10^-4^, 10^-5^, and 0) with T7 WT and Δ*0.7* in at least two biological replicates. Replicates have the same background color which alternates between white and grey. **(b)** Norm. AUC over MOI plots for strains that got infected with evidence of defense (Extended Data Fig. 10c) and were removed from the downstream effect size analysis, as we could not reproducibly validate the evidence of defense seen in the screen. Strains were treated and data are represented as described in (a). **(c)** Mean effect sizes (grey bars) and values from individual biological replicates (black circles), estimated for validated strains across biological replicates (see Methods). Outlier replicates without defense (one biological replicate for H10407 and HM-341 each, see (a)), or with nearly full defense (two biological replicates for IAI20 and IAI28 each, see (a)) were excluded from the analysis. If the lowest MOI does not allow defense against T7, but against Δ*0.7* (three biological replicates for H10407 and HM-341 each, see (a)), the effect size was estimated assuming that a 10-fold lower MOI would allow defense against T7, which may even underestimate the actual effect size.

